# Neddylation regulates the development and function of excitatory neurons

**DOI:** 10.1101/2024.12.18.629191

**Authors:** Josefa Torres, Zehra Vural, Maksims Fiosins, Fritz Benseler, Stefan Bonn, Silvio O. Rizzoli, JeongSeop Rhee, Nils Brose, Marilyn Tirard

## Abstract

The development and function of neurons is orchestrated by a plethora of regulatory mechanisms that control the abundance, localization, interactions, and function of proteins. A key role in this regard is assumed by post-translational protein modifications (PTMs). While some PTM types, such as phosphorylation or ubiquitination, have been explored comprehensively, PTMs involving ubiquitin-like modifiers (Ubls) have remained comparably enigmatic (Ubls). This is particularly true for the Ubl Nedd8 and its conjugation to proteins, i.e. neddylation, in nerve cells. In the present study, we generated a conditional Nedd8 knock-out mouse line and examined the consequences of Nedd8-deletion in cultured postmitotic glutamatergic neurons. Our findings reveal that Nedd8-ablation in young glutamatergic neurons causes alterations in the expression of developmental transcription factors that control neuronal differentiation, ultimately leading to defects in the development of a mature glutamatergic neuronal phenotype. Apparent manifestations of these defects include increased vGlut2 expression levels, reduced vGlut1 and endophilin 1 expression levels, reduced dendrite complexity, and increased transmitter release probability. Collectively, our results highlight a pivotal role for neddylation in controlling the fate of glutamatergic neurons and excitatory synaptic transmission.

**Highlights:**

- Reduced dendrite complexity in Nedd8-deficient neurons
- Increased synaptic vesicle fusion probability in Nedd8-deficient neurons
- Altered synaptic vesicle cycling in Nedd8-deficient neurons
- Neddylation controls the maturation of glutamatergic neurons

**Graphical Abstract:** **Research Topic**

## Introduction

Neurons are post-mitotic cells that usually survive for the lifespan of their host organism. To allow for such long lifetimes, the protein homeostasis of neurons is dynamically regulated and tightly controlled to allow for optimal functional efficacy and adaptability to changes in their environment. Synapses between neurons are the prime communication units in the brain. They determine the spatiotemporal organization of neuronal inputs and outputs and thereby define the layout and function of neuronal circuits, and their plasticity is a key determinant of memory processes. Accordingly, as with neurons in general, synaptic protein homeostasis and function are under tight dynamic control (Cajigas, Will, and Schuman 2010; Wang, Lauwers, and Verstreken 2017; Sun and Schuman 2022).

Post-translational protein modifications (PTM) are essential regulatory mechanisms in this regard, beyond processes that control gene transcription and mRNA translation. They expand the proteome by diversifying protein structure and function and they are at the core of many signaling pathways that enable cells to rapidly respond or adapt to changes in their physiological status and environment (Ilic, Magnussen, and Tirard 2022; Kawabe and Stegmuller 2021; Sansevrino, Hoffmann, and Milovanovic 2023; Herbst and Martin 2017; Mabb 2021). In essence, the cellular profile of PTM protein targets co-defines a specific cell-type in a defined physiological state.

The spectrum of known PTMs is broad, ranging from proteolytic protein processing and the chemical modification of amino acid side chains with various small groups to the conjugation of entire protein moieties to specific amino acids in proteins. Likewise, the functional consequences of PTMs are very diverse, affecting the interaction, function, localization, half-life, and other features of proteins (Ilic, Magnussen, and Tirard 2022; Mabb 2021; Corti and Duarte 2023). As concerns neuronal development and function, an extensive number of studies demonstrated fundamental roles of multiple PTMs, such as phosphorylation and ubiquitination, in neuronal physiology and pathophysiology (Gupta et al. 2021; Corti and Duarte 2023). Indeed, the PTM of key transcription factors orchestrates the acquisition of cell type identity, neuronal differentiation, and synapse diversity (Mihalas and Hevner 2017; Hodge, Kahoud, and Hevner 2012; Geng et al. 2021; Cui et al. 2018), and the PTM of synaptic proteins regulates every step of the synapse fate, from synaptogenesis, via synapse maturation and plasticity, to synapse maintenance and elimination (Ambrozkiewicz and Kawabe 2015; Zajicek and Yao 2021; Sansevrino, Hoffmann, and Milovanovic 2023; Mabb 2021; Moutin et al. 2021).

Based on a series of recent studies, PTMs involving ubiquitin-like modifiers (Ubls) have emerged as a novel protein regulatory principle of fundamental importance in many cell biological processes, operating at a level of complexity akin to phosphorylation or ubiquitination (Ilic, Magnussen, and Tirard 2022; Gupta et al. 2021; Mabb 2021). However, the functional consequences of PTMs by Ubls such as the SUMOs (Small Ubiquitin-like Modifiers) or Nedd8 (neuronal precursor cell expressed developmentally down regulated 8 (Kumar, Tomooka, and Noda 1992)) are still poorly understood. On the one hand, we and others showed that the sumoylation of multiple transcriptional regulators affects neuronal development and function, supporting the important notion that the PTM of key transcriptional regulators is essential for the determination of cell type identity, neuronal differentiation, and synapse diversity (Daniel et al. 2023; Ripamonti et al. 2020b; Shalizi et al. 2006; Onishi et al. 2009; Tai et al. 2016; Taylor and Labonne 2005). Conversely, the role of sumoylation in extranuclear neuronal sub-compartments, such as synapses, remains enigmatic (Daniel et al. 2017; Suk et al. 2023).

Beyond SUMOs, Nedd8 is arguably the second best characterized Ubl. Like all other Ubls, it conjugated to lysine residues in target proteins via a three-step enzymatic cascade (Kumar, Yoshida, and Noda 1993; Ilic, Magnussen, and Tirard 2022), involving an E1 (Nae1, heterodimer of Appb1 and Uba3), an E2 (Ubc12), and several E3 enzymes (RBX1/2, DCN1, MDM2, Rnf111), whereas several peptidases (Nedp1/Senp8, Uchl3, Csn5) ensure dynamics of this PTM (Ilic, Magnussen, and Tirard 2022). Neddylation is an essential PTM, and Nedd8 knock-out mice as well as mice lacking Nae1 and Ubc12 die during embryogenesis (Ilic, Magnussen, and Tirard 2022)). As with other Ubls, neddylation has mainly been studied in highly proliferating cells, where it regulates a plethora of signaling events involved in the maintenance of DNA integrity or proteostasis (Maghames et al. 2018; Lobato-Gil et al. 2021). With regard to neurons, several functional perturbation studies linked neddylation to neurite outgrowth, spine growth, synapse density, and synaptic transmission (Vogl et al. 2015; Vogl et al. 2020; Scudder and Patrick 2015; Brockmann et al. 2019; Kang et al. 2021; Song et al. 2023). However, the Nedd8 targets at the basis of these processes are unknown. The most abundant Nedd8 targets in all cell types are the cullins, components of the large multi-protein cullin-RING E3 ubiquitin ligases (Nguyen, Wang, and Xiong 2017; Petroski and Deshaies 2005) that regulate a myriad of cellular processes, including the development and dendrite arborisation of neurons (Liao et al. 2004; Ding et al. 2007; Shim et al. 2024). So far, only very few neuronal non-cullin Nedd8 targets have been described (Enchev, Schulman, and Peter 2015), including Cofilin, PSD95, and mGlu7 (Kang et al. 2021; Vogl et al. 2020; Vogl et al. 2015), whose neddylation is thought to partially account for the morphological and functional defects seen in studies on the perturbation of neddylation (Vogl et al. 2015; Vogl et al. 2020; Scudder and Patrick 2015; Brockmann et al. 2019; Kang et al. 2021; Song et al. 2023). However, a definitive and systematic understanding of the role of neddylation in neuronal development and synaptic function is lacking, not least because corresponding earlier approaches to perturb neddylation were either of unknown specificity (i.e. pharmacological inhibition of Nae1 with MLN4924) or indirect (i.e. KO of Nedd8 peptidases), making data interpretation difficult.

To obtain stringent and comprehensive insight into the role of neddylation in neurons, we generated a conditional Nedd8 mouse line and analysed the impact of Nedd8 loss on neuronal development and synaptic transmission. We systematically assessed the morphology and function of Nedd8-deficient neurons in autaptic culture, and observed defects in neuronal morphology and synaptic vesicle (SV) release probability. These changes appeared to be caused by a defect in SV cycling due to altered levels of the vesicular glutamate transporters vGlut1 and vGlut2 and of the endocytosis regulator endophilin1. Transcriptome analysis revealed that Nedd8-deficiency does not only cause strong alterations in the expression of key synapse proteins but also of transcriptional regulators involved in neuronal development and differentiation, particularly of glutamatergic neurons. In sum, our results indicate a key role for neddylation in controlling the fate of glutamatergic neurons and excitatory synaptic transmission.

## Results

Using CRISPR/Cas9, we generated a conditional Nedd8 knock-out (Nedd8cKO) mouse line by inserting LoxP sites before the second and after the third *Nedd8* exons (Figure 1A). Western blot analysis of lysates from primary hippocampal neurons infected with a CRE-expressing virus confirmed the loss of Nedd8, as compared to neuronal cultures expressing red fluorescence protein (RFP) as a control (Figure 1B). For all subsequent experiments, primary hippocampal neurons obtained from P0 Nedd8cKO pups were infected after one day *in vitro* (DIV1) with CRE-expressing virus to ablate Nedd8 expression (Nedd8-KO), RFP-expressing virus or no virus as control (CTRL), and were analyzed at DIV 9-12.

**Figure 1:**
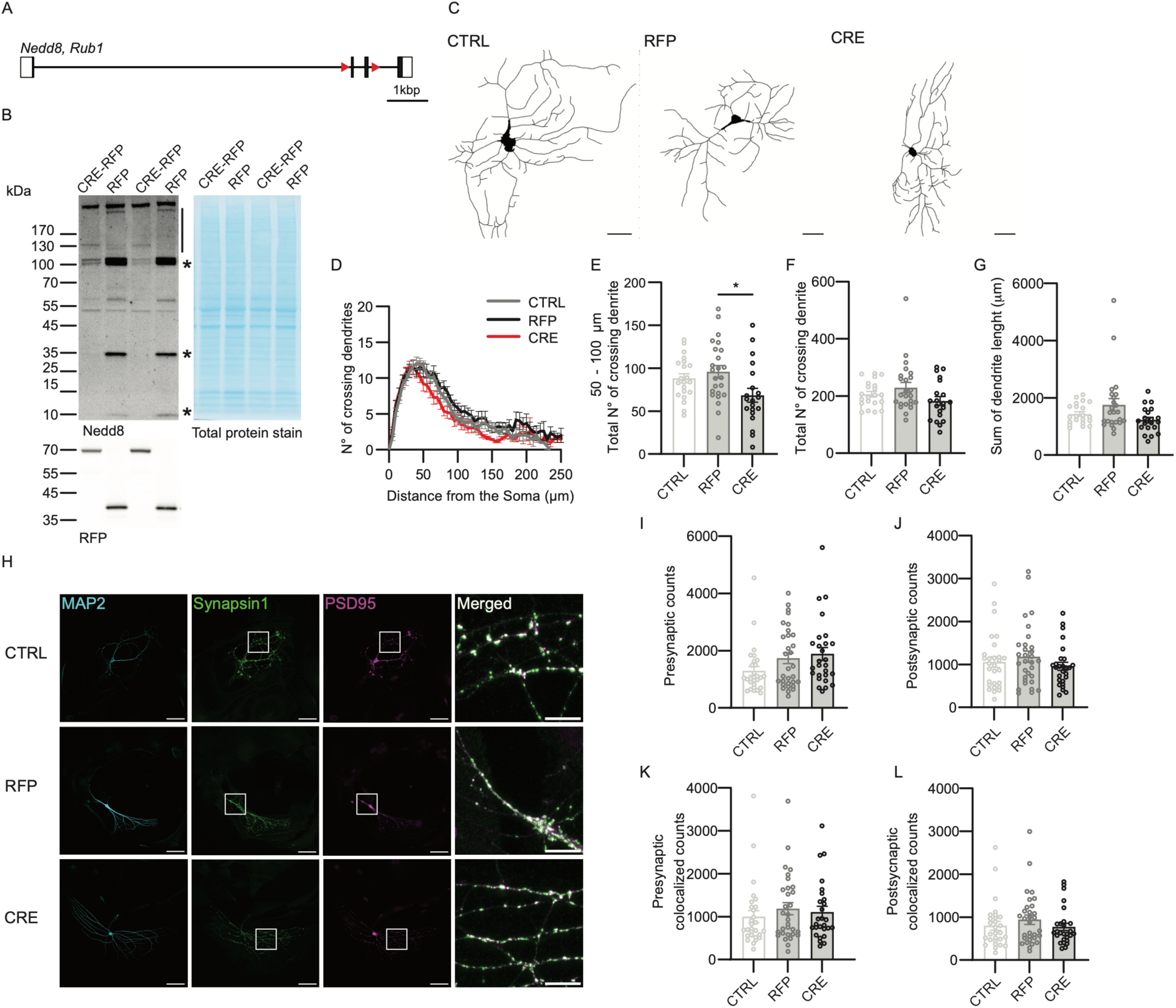
Nedd8 deficiency alters dendritic branching but not synapse number. A. On scale schematic view of the *Nedd8* mouse gene. LoxP sites (red triangles, not on scale) were inserted before the second exon and after the third exon using a CRISPR/Cas9 strategy. B. Anti-Nedd8 (top) and RFP (bottom) Western blot analysis of DIV12 primary hippocampal culture lysates induced with virus as indicated on top. The line and stars indicate NEdd8-specific signal. Molecular weight marker is indicated on the left (kDa). C. DIV13 Nedd8cKO autaptic hippocampal neurons infected with lentivirus expressing RFP or CRE-recombinase-RFP (CRE) were fixed and immunostained for MAP2. Neurons without lentiviral infection were used as infection controls (CTRL). Images were acquired using a confocal microscope and representative binary images of MAP2-immunostained neurons are shown. Scale bar = 20 μm. D. Line plots showing the number of dendritic intersections as a function of distance from the soma obtained by Sholl analysis (N=3; n_CTRL_=22, n_RFP_=26, n_CRE_=20 cells). E. Bar graph showing the number of crossing dendrites between 50 and 100 μm from the soma, calculated from Sholl analysis (N=3; n_CTRL_=22, n_RFP_=26, n_CRE_=20 cells). F. Bar graphs showing the total number of crossing dendrites, calculated from Sholl analysis (N=3; n_CTRL_=22, n_RFP_=26, n_CRE_=20 cells). G. Bar graph showing the dendrite length sum (N=3; n_CTRL_=22, n_RFP_=26, n_CRE_=20 cells). H. Immunostaining of autaptic hippocampal neurons with antibodies against MAP2 (cyan), synapsin1 (green), and PSD95 (magenta). Scale bar = 50 μm. The white square indicates a closer view depicted in the right column of panels. Scale bar = 10 μm. I. Bar graph showing the total number of synapsin1 (presynaptic) counts. (N=3; n_CTRL_=28, n_RFP_=31, n_CRE_=27 cells). JI. Bar graph showing the total number of PSD95 (postsynaptic) counts (N=3; n_CTRL_=28, n_RFP_=31, n_CRE_=27 cells). K. Bar graph showing the total number of counts of synapsin1 colocalized with PSD95 signal (N=3; n_CTRL_=28, n_RFP_=31, n_CRE_=27 cells). LK. Bar graphs showing the total number of counts of PSD95 colocalized with synapsin1 (N=3; n_CTRL_=28, n_RFP_=31, n_CRE_=27 cells). In each graph, dots correspond to the total number of synaptic counts obtained from one individual neurons. Bars represent mean ±SEM. Data was compared using a Kruskal-Wallis and Dunńs multiple comparison test, where * p<0.05.

### Nedd8-KO neurons show morphological defects but no change in synapse number

Many studies described defects in neuronal development, morphology and synapse number upon neddylation blockade (Song et al. 2023; Vogl et al. 2015; Kang et al. 2021). Thus, we analysed these parameters in Nedd8-deficient neurons as compared to RFP or non-infected control neurons (Figure 1C-L). Sholl analysis revealed that the complexity of the neuronal dendritic tree was slightly altered upon Nedd8-deficiency, with a minor, but significant, reduction in the number of crossing dendrites within 50-100 μM from the soma (Figure 1C-E). However, the total number of crossing dendrites and the total dendrite length were not reduced (Figure 1E-G), indicating a moderate, but significant, impact of Nedd8 depletion on neuronal morphology.

Next, we quantified the number of excitatory synapses via immunolabeling using Synpasin1 as pre-synaptic marker and PSD95 to label excitatory post-synapses (Figure 1H-L); MAP2 was used to visualise overall neuronal processes. The number of pre- and post-synaptic puncta was similar in Nedd8-KO as compared to the controls, e.g. RFP or non-infected CTRL neurons (Figure 1I, J), leading to similar number of pre- and postsynaptic colocalized puncta (Figure 1K, L). Taken together, our data indicate that loss of Nedd8 expression alters neuronal morphology only moderately, without any impact on the number of synapses.

### Altered synaptic transmission in Nedd8-KO neurons

Next, we examined how Nedd8 depletion affected glutamatergic neurotransmission in autaptic hippocampal neurons (Figure 2 and 3). While Nedd8-deficient autaptic neurons displayed no change in evoked excitatory post-synaptic currents (EPSCs) as compared to RFP or non-virus infected neurons (CTRL, Figure 2A), CRE-infected neurons showed a 30% decrease in the readily-releasable pool (RRP) of synaptic vesicles as assessed by hypertonic sucrose stimulation (Figure 2B), accompanied with a significant increase in synaptic transmitter release probability (P_vr_, Figure 2C). Importantly, there was no change in spontaneous miniature EPSC (mEPSC) amplitude (Figure 2D), while a non-significant trend towards a reduction in mEPSC frequency was observed (Figure 2E). Importantly, currents induced by fast superfusion of neurons with glutamate or GABA were not different between Nedd8-KO and the CTRL and RFP controls (Figure 2F, G), indicating no differences in the surface expression of functional glutamate receptors. Thus, Nedd8 depletion does not profoundly alter post-synaptic function in a profound fashion.

**Figure 2:**
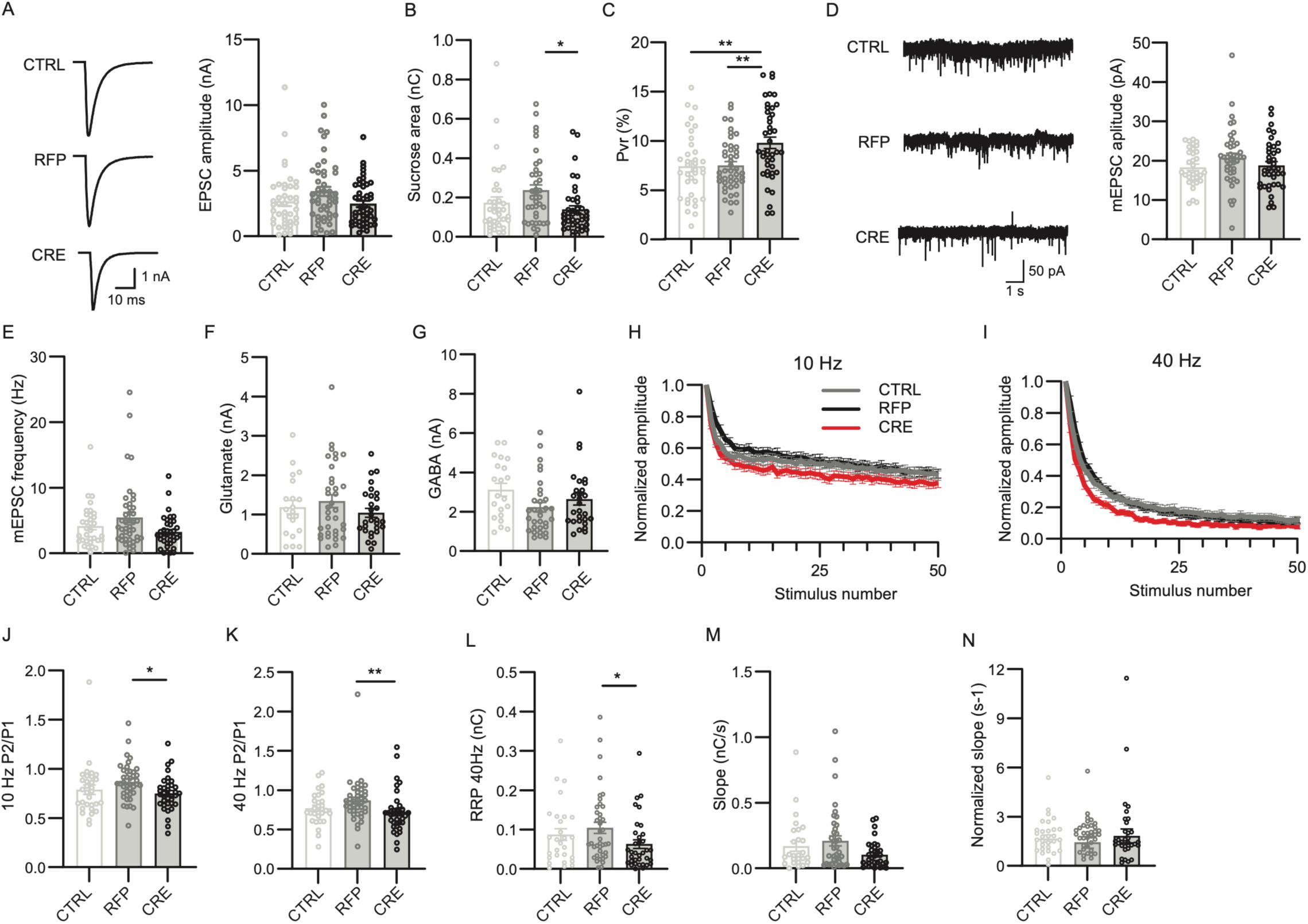
Nedd8 knock-down alters synaptic release probability. Autaptic excitatory hippocampal neurons from Nedd8cKO animals were infected with lentivirus expressing RFP or CRE-recombinase-RFP (CRE). Neurons without lentiviral infection were used as infection controls (CTRL). A. Representative traces (left) and bar graph (right) of evoked EPSC amplitudes of autaptic neurons infected as indicated below the graphs (N=5; n_CTRL_=41, n_RFP_=47, n_CRE_=47 cells). B. Bar graph depicting the total charge transferred by the release of the RRP obtained after treatment with hypertonic sucrose solution (N=5; n_CTRL_=36, n_RFP_=44, n_CRE_= 42 cells). C. Bar graph showing the probability of vesicular release (P_vr_) of individual neurons (N=5; n_CTRL_=36, n_RFP_=44, n_CRE_= 42 cells). P_vr_ is calculated by dividing the charge transfer during an evoked EPSC by the charge transfer during the sucrose response, and then expressed as a percentage. D. Representative traces (left) and bar graph (ritght) depicting spontaneous mEPSC amplitudes of autaptic neurons treated with 300nM TTX (N=5; n_CTRL_=33, n_RFP_=43, n_CRE_= 39 cells). E. Bar graph showing the frequency of mEPSC measured in autaptic neurons in the presence of 300nM TTX (N=5; n_CTRL_=33, n_RFP_=43, n_CRE_= 39 cells). P between RFP and CRE = 0.0914 F. Bar graph showing the amplitude of the peak current generated by the superfusion of 100 μM Glutamate (N=5; n_CTRL_=20, n_RFP_=34, n_CRE_= 29 cells). G. Bar graph showing the amplitude of the peak current generated by the superfusion of 3 μM GABA (N=5; n_CTRL_=21, n_RFP_=32, n_CRE_= 28 cells). H. Line plots showing the change in EPSC amplitudes during a 10 Hz stimulation train (N=5; n_CTRL_=31, n_RFP_=41, n_CRE_= 37 cells). Data were normalised to the first response of the respective train and shown as mean +/- SEM. I. Line plots showing the change in EPSC amplitudes during a 40 Hz stimulation train (N=5; n_CTRL_=29, n_RFP_=42, n_CRE_= 37 cells). Data were normalised to the first response of the respective train and shown as mean +/- SEM. J. Bar graph showing paired-pulse ratios obtained from 10 Hz stimulation trains (N=5; n_CTRL_=31, n_RFP_=41, n_CRE_= 37 cells). K. Bar graph showing paired-pulse ratios obtained from 40 Hz stimulation trains (N=5; n_CTRL_=29, n_RFP_=42, n_CRE_= 37 cells). L. Bar graphs showing the RRP size as estimated by back extrapolation of the cumulative EPSC after 40 Hz stimulus trains (N=5; n_CTRL_=28, n_RFP_=40, n_CRE_= 35 cells). M. Bar graphs showing the slope of the back extrapolated cumulative EPSC curve of the 40 Hz trains (N=5; n_CTRL_=28, n_RFP_=39, n_CRE_= 35 cells). N. Bar graph showing the ratio between the slope of the back extrapolated cumulative EPSC curve (shown in M), divided by the RRP_40Hz_ (shown in L), (N=5, n_ctrl_ = 32, n_rfp_ = 39, n_cre_ = 34 cells). In each graph, dots correspond to measurements recorded from individual neurons and bars represent mean ±SEM. Data was compared using a Kruskal-Wallis and Dunńs multiple comparison test, where *p<0.05, **p<0.01.

**Figure 3:**
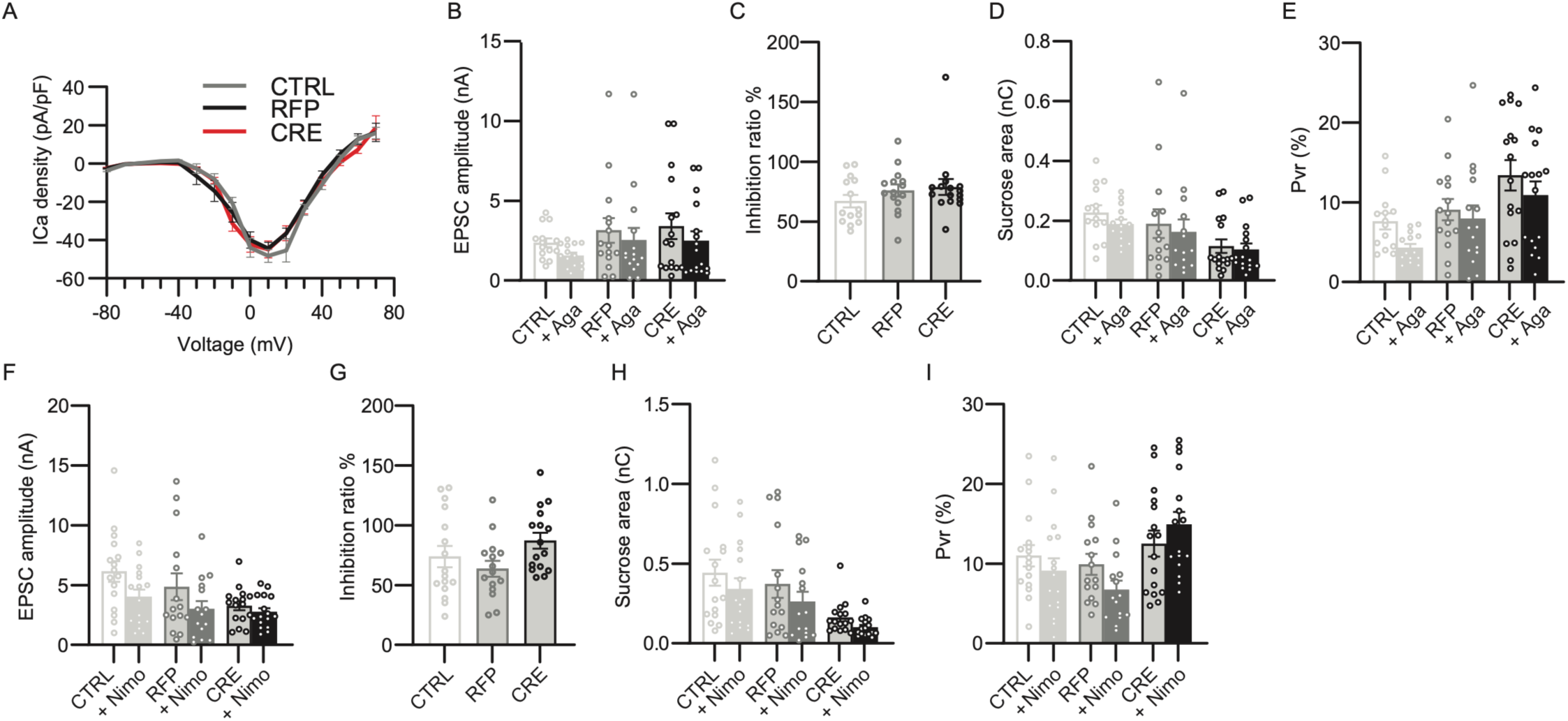
VGCC currents are unchanged in Nedd8-KO neurons. Analysis of calcium currents obtained from DIV9-13 Nedd8cKO autaptic excitatory hippocampal neurons infected with lentivirus expressing RFP, CRE-recombinase-RFP (CRE) or non-infected neurons (CTRL). A. Current-voltage relationship of HVA calcium channels (N=3; n_CTRL_=23, n_RFP_=21, n_CRE_=23 cells). B. Bar graph showing the change in evoked EPSC amplitudes after treatment with 0.2 µM ω-Agatoxin (Aga) (N=2; n_CTRL_=14, n_RFP_=15, n_CRE_=16 cells). C. Bar graph showing the inhibition ratio of EPSC amplitudes after treatment with 0.2 µM ω-Agatoxin (N=2; n_CTRL_=14, n_RFP_=15, n_CRE_=16 cells). D. Bar graph depicting the total charge transferred by the release of the RRP upon application with hypertonic sucrose solution after treatment with 0.2 µM ω-Agatoxin (Aga) (N=2; n_CTRL_=14, n_RFP_=15, n_CRE_=16 cells). E. Bar graph showing the probability of vesicular release of individual neuron after treatment with 0.2 µM ω-Agatoxin (Aga) (N=2; n_CTRL_=14, n_RFP_=14, n_CRE_=16 cells). F. Bar graph showing the change in EPSC amplitudes after treatment with 10 µM Nimodipine (Nimo) (N=2; n_CTRL_=16, n_RFP_=15, n_CRE_=16 cells). G. Bar graph showing the inhibition ratio of EPSC amplitudes after treatment with 10 µM Nimodipine (N=2; n_CTRL_=16, n_RFP_=15, n_CRE_=16 cells). H. Bar graph of the total charge transferred by the release of the RRP upon application with hypertonic sucrose solution after treatment with 10 µM Nimodipine (Nimo) (N=2; n_CTRL_=16, n_RFP_=15, n_CRE_=16 cells). I. Bar graph showing the probability of vesicular release of individual neurons after treatment with 10 µM Nimodipine (Nimo) (N=2; n_CTRL_=16, n_RFP_=15, n_CRE_=16 cells). In each graph, dots correspond to measurements recorded from individual neurons and bars represent mean ±SEM. Data was compared using unpaired *t*-test, Mann-Whitney test, and a Kruskal-Wallis and Dunńs multiple comparison test.

We next studied synaptic short-term plasticity by measuring the synaptic response to high frequency stimulation (HFS) (Figure 2H-N). We found that the EPSC amplitude of CRE-infected neurons depressed faster than that of RFP or non-infected CTRL neurons (Figure 2H, I). Additionally, and in accord with the decreased RRP size as assessed by hypertonic sucrose stimulation (Figure 2B), the paired pulse ratio (Figure 2J, K) and the RRP size (RRP_40Hz_) as estimated by back extrapolation of the cumulative EPSC during 40Hz stimulus trains were decreased in Nedd8-deficient neurons as compared to RFP or non-infected CTRL neurons (Figure 2L). Finally, the refiling rate of the RRP – i.e. the slope of the back extrapolated cumulative EPSC curve of the 40 Hz trains – showed a trend towards reduction that was not significant (Figure 2M). When we normalized the RRP refiling rate to the RRP_40Hz,_ no difference between the tested group were detectable (Figure 2N). Thus, the more pronounced short-term depression observed in Nedd8-deficient neurons (Figure 2H) and 2I) is primarily attributable to a reduction in RRP size and increase P_vr_, rather than to an aberrant activity-dependent RRP refiling process.

### Normal VGCC activity but reduced SV-VGCC coupling in Nedd8-KO neurons

Regarding the changes in short-term plasticity observed in Nedd8-deficient neurons, we next examined the possibility of alterations in the properties of voltage-gated Ca^2+^ channels (VGCCs). Given the serious challenges associated with studying VGCCs in the small pre-synaptic terminals of cultured hippocampal neurons, we investigated the VGCC properties in the soma of proximal dendrites. Initially, we assessed the overall density and voltage-dependence of VGCCs by measuring the current-voltage relationships, and found that Nedd8-loss had no discernible effect (Figure 3A). We expanded our analyses by employing VGCC subtype-specific blockers to determine whether the influence of different presynaptic VGCCs in the presynaptic terminal on EPSC amplitude, RRP size, and P_vr_ is altered by Nedd8-loss, by administering ω-Agatoxin and nimodipine, which specifically inhibit P/Q-type and L-type VGCCs, respectively (Figure 3B-3I). We found no significant differences between the groups tested, indicating that the altered RRP and P_vr_ in Nedd8-KO neurons cannot be attributed to modification in VGCCs properties.

In light of the results outlined above, particularly the P_vr_ increase whithout changes in VGCC function, we next explored the possibility that Nedd8-loss selectively affects the priming and fusion of SVs that are more readily available for synaptic transmitter release due, to VGCC-coupling. We performed electrophysiological recordings in the presence of EGTA-AM, a slow Ca^2+^ chelator used to perturb VGCC coupling to SV Ca^2+^ sensor (Nakamura 2019; Ritzau-Jost et al. 2018) (Figure 4A). EGTA-AM reduced the P_vr_ to a similar extent in Nedd8-KO neurons as compared to RFP infected neurons (Figures 4B-4D). Strikingly, however, EGTA-AM reversed synaptic depression during trains of action potentials in RFP-infected neurons (Figure 4E and 4G) but not in CRE-expressing Nedd8-deficient cells (Figure 4F-4H). These data are compatible with the notion that a pool of ready-releasable SVs present near VGCCs is altered in Nedd8-deficient neurons (Figure 4A).

**Figure 4:**
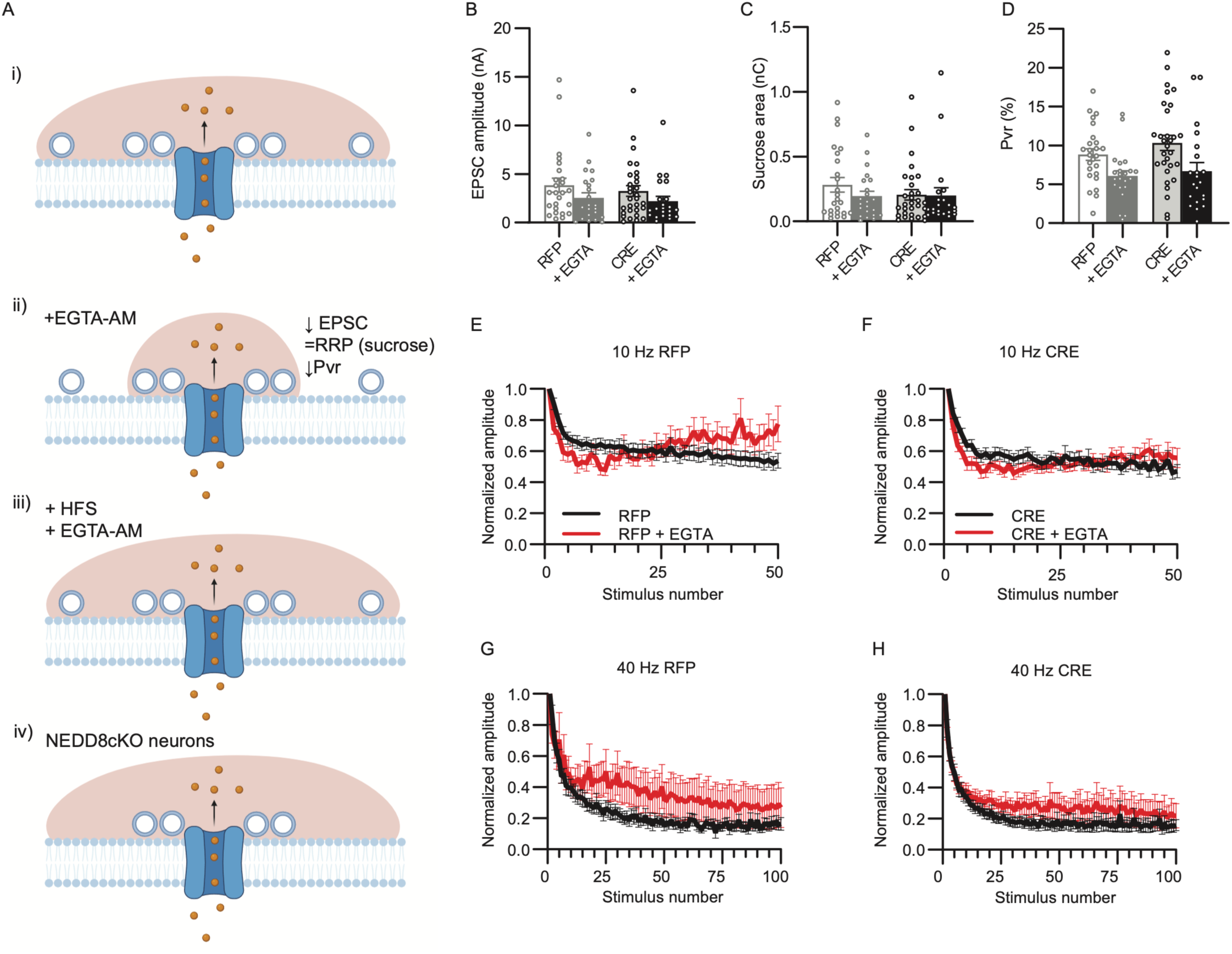
Nedd8 regulates the number of synaptic vesicles coupled to VGCC. Recordings were performed using DIV9-13 Nedd8cKO autaptic excitatory hippocampal neurons infected with lentivirus expressing RFP, CRE-recombinase-RFP (CRE) or non-infected neurons (CTRL). A. Scheme representing the effect of EGTA-AM on synaptic vesicles coupled to calcium channels. i) Following an action potential, voltage-gated calcium channels (VGCC) open and Ca^2+^ ions access the intracellular compartment. The red cloud indicates the area and number of vesicles impacted by the entry of calcium. ii) After treatment with the permeable ester, EGTA-AM, hydrolyzation of intracellular estearses leads to the production of EGTA, which lowers the intracellular calcium concentration (reduction in the size of the red cloud), except in the vicinity of VGCC, leading to a decrease in the number of released vesicles. Ultimately, EPSC amplitude and P_vr_ are reduced, but not the size of the RRP, as measured by hypertonic sucrose treatment. iii) During a high frequencies train of action potentials (HFS), the effect of EGTA is restored due to accumulation of calcium ions in the intracellular compartment. iv) Working model: Nedd8-deficient neurons present fewer synaptic vesicles near VGCC. B. Bar graph showing EPSC amplitudes, with and without treatment with 100 µM EGTA-AM (N=3; n_RFP_=14, n_RFP+EGTA-AM_=16, n_CRE_=20, n_CRE+EGTA-AM_=15 cells). C. Bar graph of the total charge transferred by the release of the RRP upon application of hypertonic sucrose solution, with and without treatment with 100 µM EGTA-AM (N=3; nRFP=14, nRFP+EGTA-AM=17, nCRE=19, nCRE+EGTA-AM=15 cells). D. Bar graph showing the impact of EGTA-AM treatment on the probability of vesicular release of individual neuron (N=3; n_RFP_=14, n_RFP+EGTA-AM_=17, n_CRE_=20, n_CRE+EGTA-AM_=15 cells). E. Line plots showing the change in EPSC amplitudes during a 10 Hz stimulation train in neurons infected with RFP virus, with and without application of 100 µM EGTA-AM (N=3; n_RFP_=15, n_RFP+EGTA-AM_=16 cells). F. Line plots showing the change in EPSC amplitudes during a 10 Hz stimulation train in neurons infected with CRE virus, with and without application of 100 µM EGTA-AM (N=3; n_CRE_=19, n_CRE+EGTA-AM_=14 cells). G. Line plots showing the change in EPSC amplitudes during a 40 Hz stimulation train in neurons infected with RFP virus, with and without application of 100 µM EGTA-AM (N=3; n_RFP_=15, n_RFP+EGTA-AM_=16 cells). H. Line plots showing the change in EPSC amplitudes during a 40 Hz stimulation train in neurons infected with CRE virus, with and without application of 100 µM EGTA-AM (N=3; n_CRE_=19, n_CRE+EGTA-AM_=14 cells). In each graph, dots correspond to measurements recorded from individual neurons and bars represent mean ±SEM. Data was compared using an unpaired t-test and a Mann-Whitney test. Illustrations were made using Briorender.com.

### Altered expression of genes encoding key pre-synaptic proteins in Nedd8-KO neurons

In a first attempt to explore the molecular pathways by which Nedd8-loss causes the functional phenotypes we observed, we performed RNA-sequencing and differential expression analysis of CRE-expressing neurons vs. RFP-expressing neurons (Figure 5A, Supplementary Figures 1-3, Table 1). The underlying idea was to not only determine direct transcriptional changes upon Nedd8-loss, but to also assess indirect compensatory mechanisms. Accordingly, we put one focus on synaptic components using the SynGO database (Koopmans et al. 2019).

**Figure 5:**
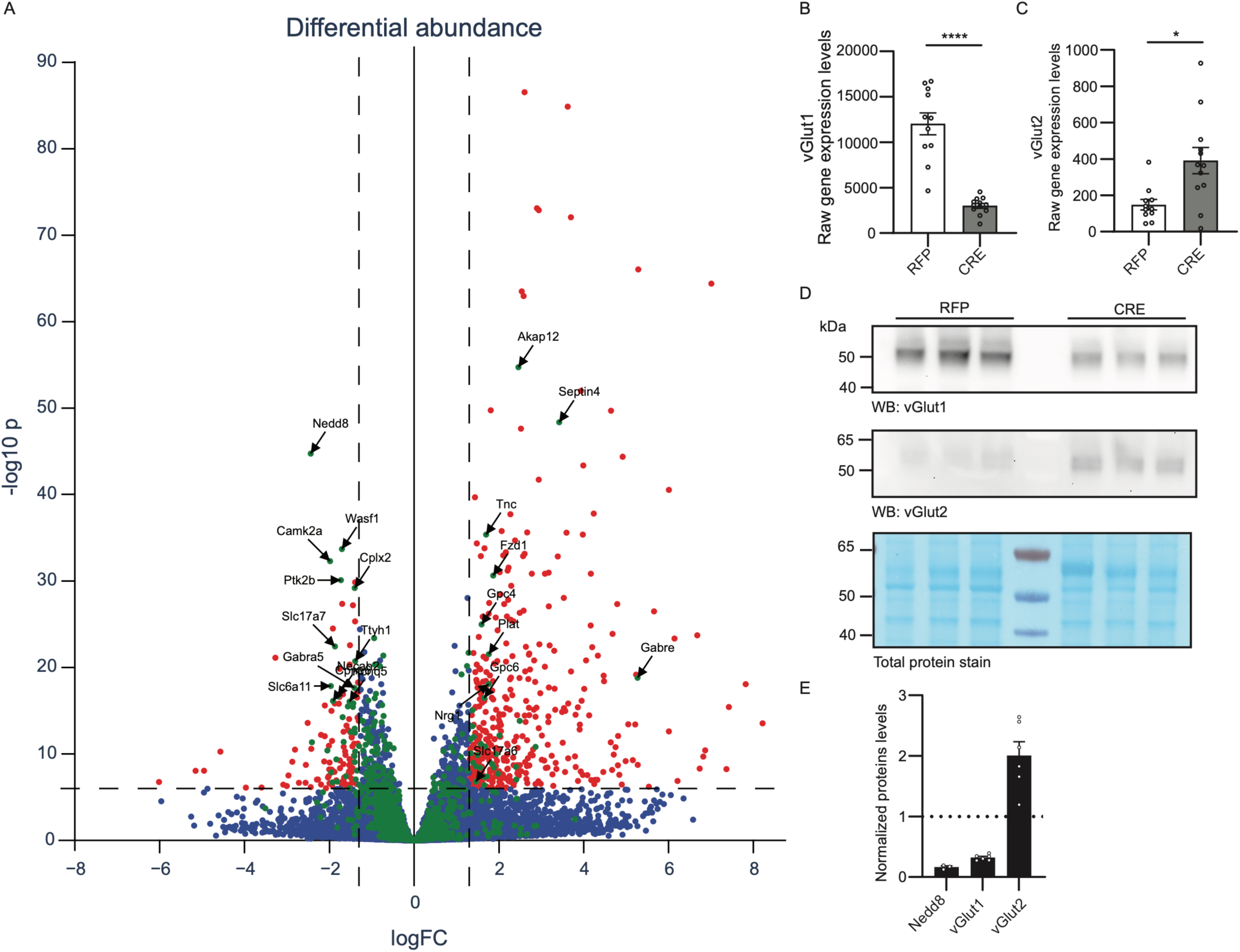
Nedd8 depletion alters the expression of synaptic proteins. A. RNA was extracted from DIV13 Nedd8cKO primary hippocampal neurons infected with lentivirus expressing RFP or CRE-recombinase-RFP (CRE) and sequenced. Each dote represent a single gene (blue dots). Volcano plot depict each gene based on their Log2 fold change (LogFC) and the reverse log10 of their p value (-log10p). Red dots correspond to genes with a LogFC above 1,3 and -log10p above 6. Green dots correspond to genes found in SynGO. B. Bar graphs showing the raw gene expression levels of vGlut1 (N=3, n_RFP_=11, n_CRE_=12). C. Bar graphs showing the raw gene expression levels of vGlut2 (N=3, n_RFP_=11, n_CRE_=12). D. Total protein stain (bottom panel) and anti-vGlut1 and vGlut2 Western blot analysis of primary hippocampal Nedd8cKO neurons lysates infected with lentivirus expressing RFP or CRE-recombinase-RFP (CRE). Molecular weight is indicated on the left side (kDa). E. Bar graph depicting normalised protein levels of Nedd8, vGlut1 and vGlut2, as assessed by Western blot in D.

**Table 1:** List of differentially expressed genes (DE), list of differentially expressed genes after filtering (DE_filtered), list of differentially expressed genes found in the SynGO database (DE_filtered_syngo).

Upon stringent filtering, the expression of 566 genes remained significantly altered in Nedd8-deficient neurons (Figure 5A; Log2FC of 1.3; -log10 p-value of 6), with 77% (436) up- and 23% (130) down-regulated. Among these were 66 SynGO genes that remained significantly changed after filtering. Interestingly, half of these (33) were up- (33) and the other half (33) down-regulated (Figure 5A), which strongly contrasts with the remodeling of the rest of the proteome induced by Nedd8-deficiency, indicating that Nedd8-loss causes a general increase in gene expression but a more pronounced downregulation of genes expressing synaptic proteins.

SynGO pathway analysis showed that Nedd8-deletion leads to decreased expression of multiple mRNAs encoding pre-synaptic components, in particular proteins involved in the SV cycle (Supplementary Figure 2) and to the up-regulation of genes related to the extracellular matrix or protein translation (Supplementary Figure 3). Strikingly, vGlut1 (Slc17a7) was strongly down-regulated upon Nedd8-loss while vGlut2 was up-regulated (Slc17a6, Figure 5A, B). This result was further validated at the protein level (Figures 5C). As there is a correlation between the vGlut isoform expressed and the P_vr_, with vGlut2-expressing neurons exhibiting higher P_vr_ (Wojcik et al. 2004; Weston et al. 2011), we further studied this phenomenon.

### Reduced density of vGlut1-containing synapses in Nedd8-KO neurons

We next assessed vGlut1 immunosignals in Nedd8-deficient neurons as compared to RFP-expressing controls (Figure 6). While the total number of synapses was unchanged (Figures 1H-L), the total number of vGlut1 puncta was significantly reduced in Nedd8-deficient neurons (Figures 6A-C). Importantly, vGlut1 signal intensity was not altered in vGlut1-containing mutant synapses (Figures 6D, 6E). STED imaging confirmed that the intensity of vGlut1 puncta along dendrites was unchanged and showed further that the sub-synaptic distribution of vGlut1 is similar between Nedd8-deficient and control neurons (Figures 6F-H). Our data indicate that the number of synapses containing robust levels of vGlut1 is decreased in Nedd8-KO neurons while the synaptic enrichment of vGlut1 is unchanged in synapses that still contain vGlut1.

**Figure 6:**
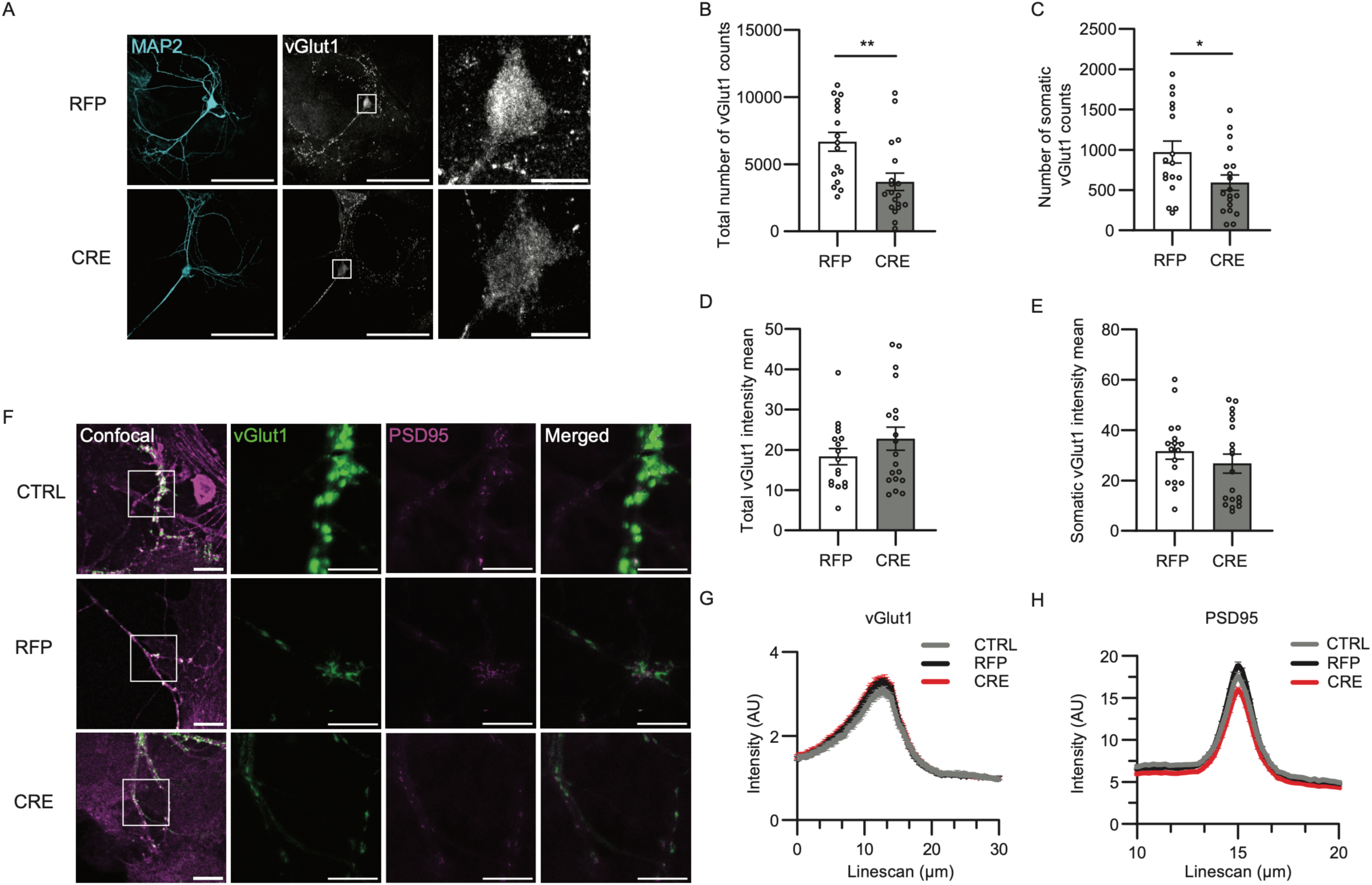
vGlut1 levels are altered upon Nedd8 depletion. DIV13 Nedd8cKO autaptic hippocampal neurons infected with lentivirus expressing RFP or CRE-recombinase-RFP (CRE) were fixed and immunostained with antibodies as indicated. Neurons without lentiviral infection were used as infection controls (CTRL). A. Representative confocal images of autaptic neurons stained with antibodies against MAP2 and vGlut1. Scale bars = 80 and 10 μm. B. Bar graph showing the total number of vGlut1 counts (N=2; n_RFP_=17, n_CRE_=19 cells). C. Bar graph showing the number of somatic vGlut1 counts (N=2; n_RFP_=17, n_CRE_=19 cells). D. Bar graph showing the total vGlut1 intensity mean (N=2; n_RFP_=17, n_CRE_=19 cells). E. Bar graph showing the somatic vGlut1 intensity mean (N=2; n_RFP_=17, n_CRE_=19 cells). F. Representative confocal (left column of panels) and STED images of autaptic hippocampal neurons labelled with nanobodies against vGlut1 (green) and PSD95 (magenta). Scale bars = 10 μm (left column of panels) and 5 μm (other panels). G. Line plots showing the change in vGlut1 signal intensity against distance (N=3; n_CTRL_=13, n_RFP_=10, n_CRE_=13 cells). H. Line plots showing the change in PSD95 signal intensity against distance (N=3; n_CTRL_=13, n_RFP_=10, n_CRE_=13 cells). In each graph, dots correspond to the total number of synaptic counts obtained from one individual neurons. Bars represent mean ±SEM. Data was compared using a Kruskal-Wallis and Dunńs multiple comparison test or Mann-Whitney test, where *p<0.05, **p<0.01

### P**_vr_** defects in Nedd8-KO neurons are not reverted by vGlut2 knock-down or vGlut1 overexpression

Our data show that vGlut2 expression is up-regulated by 50% and vGlut1 down-regulated by 60-70% upon Nedd8-loss (Figure 5), which could indicate a delayed differentiation of glutamatergic neurons, which switch from vGlut2 to vGlut1 expression as they mature, or a complex scenario of vGlut1 depletion compensated by vGlut2 expression (Boulland et al. 2009). Previous studies showned that the vGlut1-vGlut2 ratio co-defines the synaptic transmitter release probability, with vGlut2-expressing neurons being associated with higher release probability (Weston et al. 2011), an observation that could (partly) explain the release probability phenotype of Nedd8-KO neurons (Figure 2). To test this possibility, we performed rescue experiments and either re-expressed vGlut1 (Figure 7A) or knocked-down vGlut2 (Figure 7B) in the Nedd8-KO background in order to reinstall a normal vGlut1-vGlut2 ratio, and studied the impact on the RRP and P_vr_. However, neither overexpression of vGlut1 nor knock-down of vGlut2 rescued the RRP and P_vr_ phenotype of Nedd8-KO neurons (Figures 7C-H). Crucially, endophilin1 levels were also shown to correlate with the reduced release probability associated with vGlut1 expression in neurons (Kang et al. 2021). Strikingly, Nedd8-KO causes a 50% reduction in endophilin1 expression, both at the mRNA and protein levels, and re-expression of vGlut1 or knock-down of vGlut2 did not rescue endophilin1 levels in Nedd8-KO cells (Figures 7I-M). Thus, it is possible that altering vGlut1 or vGlut2 levels could not rescue the synaptic defects in Nedd8-KO neurons because of decrease in endophilin 1 levels, which cannot be compensated by changing vGlut1-vGlut2 ratios.

**Figure 7:**
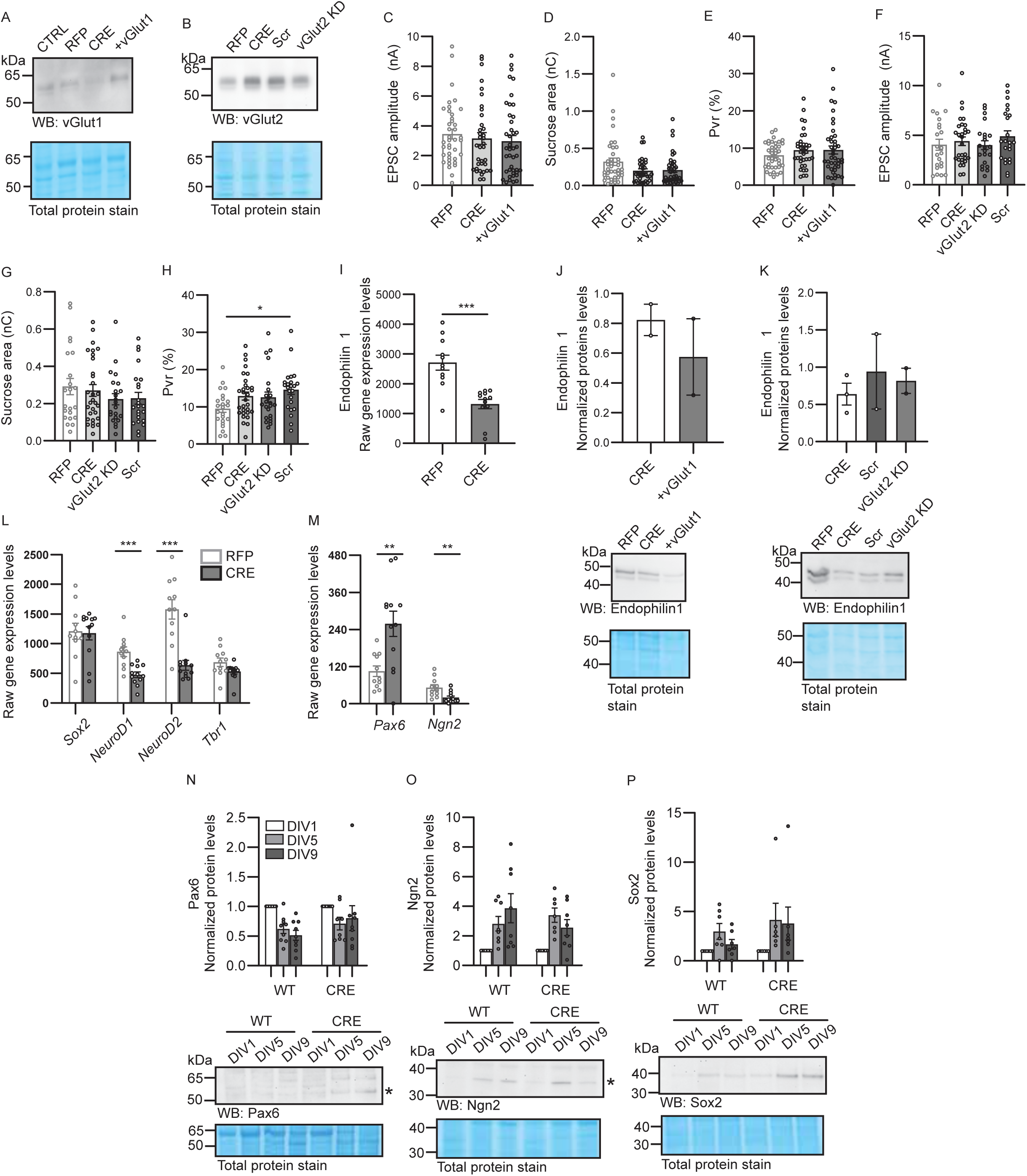
Over expression of vGlut1, or knock-down of vGlut2 do not rescue the defects in release probability. Hippocampal neurons from Nedd8cKO animals were infected with lentivirus expressing RFP, CRE-recombinase-RFP (CRE), co-infection with CRE and a virus expressing vGlut1 (+vGLUT1), co-infection with CRE and and a virus expressing shRNA to knock-down the expression of vGlut2 (vGLUT2 KD), or co-infection with CRE and a virus expressing scrambled RNA sequences (Scr). Neurons were lysed on DIV12. A-B. Total protein stain (bottom panels), anti-vGlutt1 (A) and anti-vGlut2 (B) Western blot analysis of Nedd8cKO primary hippocampal neurons lysates infected as described above. Molecular weight is indicated on the left side (kDa). C. Bar graph showing evoked EPSC amplitudes of autaptic excitatory neurons infected with lentivirus expressing RFP, CRE-recombinase-RFP (CRE), or CRE-recombinase-RFP (CRE) and a virus re-rexpressing vGlut1 (+vGlut1) (N=2, n_RFP_=32, n_CRE_=26, n_vGlut1_=27). D. Bar graph showing the total charge transferred by the release of the RRP upon application of a hypertonic sucrose solution in autaptic excitatory neurons infected with lentivirus expressing RFP, CRE-recombinase-RFP (CRE), or CRE-recombinase-RFP (CRE) and a virus re-rexpressing vGlut1 (+vGlut1) (N=2, n_RFP_=32, n_CRE_=27, n_vGlut1_=28). E. Bar graph showing the probability of vesicular release of autaptic excitatory neurons infected with lentivirus expressing RFP, CRE-recombinase-RFP (CRE), or CRE-recombinase-RFP (CRE) and a virus re-rexpressing vGlut1 (+vGlut1) (N=2, n_RFP_=32, n_CRE_=27, n_vGlut1_=28). F. Bar graph showing evoked EPSC amplitudes of autaptic excitatory neurons infected with lentivirus expressing RFP, CRE-recombinase-RFP (CRE), CRE-recombinase-RFP and a virus to knock-down the expression of vGlut2 (vGlut2 KD) or with a control virus (Scr) (N=3, n_RFP_=15, n_CRE_=19, n_vGlut2_=16, n_scramble_=15). G. Bar grap of the total charge transferred by the release of the RRP upon application of a hypertonic sucrose solution in autaptic excitatory neurons infected with lentivirus expressing RFP, CRE-recombinase-RFP (CRE), CRE-recombinase-RFP and a virus to knockdown the expression of vGlut2 (vGlut2 KD) or with a control virus (Scr) (N=3, n_RFP_=15, nCRE=19, nvGlut2=16, nscramble=15). H. Bar graph showing the probability of vesicular release in autaptic excitatory neurons infected with lentivirus expressing RFP, CRE-recombinase-RFP (CRE), CRE-recombinase-RFP and a virus to knock-down the expression of vGlut2 (vGlut2 KD) or with a control virus (Scr) (N=3, n_RFP_=15, n_CRE_=19, n_vGlut2_=16, n_scramble_=16). I. Bar graphs showing the raw gene expression levels of Endophilin1 (N=3, n_RFP_=11, n_CRE_=12). J-K. Anti-endophilin1 Western blot analysis of Nedd8cKO primary hippocampal neurons lysates not infected (CTRL) or infected with viruses expressing RFP, CRE-recombinase-RFP (CRE), CRE-recombinase-RFP (CRE) and a virus re-rexpressing vGlut1 (+vGlut1), CRE-recombinase-RFP and a virus to knock-down the expression of vGlut2 (vGlut2 KD) or with a control virus (Scr) (vGlut2 KD, Scr, K). Molecular weight is indicated on the left side (kDa). L-M. Normalised expression levels of Endophilin1 as assessed by Western blot in neurons re-rexpressing vGlut1(+vGLUT1, L) or knock-downed for vGlut2 (vGlut2 KD) as compared to control (Scr or RFP). N-O. Bar graph showing the raw gene expression levels of different transcription factors as indicated below the graph (N=3, n_RFP_=11, n_CRE_=12). Panel C to F: In each graph, dots correspond to measurements recorded from individual neurons. Panel I, L to O: in each graph, dot represent measurement obtained from individual animals. Bar represent mean ±SEM. Data was compared using a Kruskal-Wallis and Dunńs multiple comparison test, Mann-Whitney or unpaired t-test, where *p<0.05, **p<0.01, ***p<0.001, ****p<0.0001.

### Neddylation regulates excitatory neurodevelopment

We show that Nedd8 ablation leads to minor morphological changes but strong alterations in presynaptic transmitter release probability and the cycling of synaptic vesicles. To explore the biological processes underlying these morphological and functional defects, we examined changes in the neuronal transcriptional landscape of mature (DIV12) neurons upon Nedd8 deletion (Figure 5). Our transcriptome analysis has revealed that Nedd8-KO alters the expression of genes encoding synaptic proteins involved in the release and cycling of synaptic vesicles, consistent with the functional deficits we observed. These phenotypes are likely to be the culmination of a more profound, as yet unknown, altered Nedd8-mediated signaling earlier in neuronal development. Given that the most abundant Nedd8 targets are the cullins, which are large multiprotein cullin-RING E3 ubiquitin ligases (Nguyen, Wang, and Xiong 2017; Petroski and Deshaies 2005) that regulate a variety of cellular processes, including neuronal development and dendritic arborization of neurons (Liao et al., 2004, Ding et al., 2007, Shim et al., 2024), we hypothesized that the lack of degradation of key regulatory factors was at the origin of the transcriptional reshaping and functional changes we observed. For instance, our transcriptome analysis of developed neurons (DIV12) revealed a strong alteration in the expression profile of transcriptional regulators that are essential for the development and maturation of the glutamatergic neuron phenotype, e.g. *Sox2*, *NeuroD1*, *NeuroD2*, *Tbr1*, *Pax6* or *Neurogenin2* (Figure 7L, M). Thus, it is likely that Nedd8-depletion alters, directly or indirectly, the protein expression levels of such regulators during the course of neuronal maturation. To test our hypothesis, we further evaluated the protein expression levels of selected regulators during the maturation of primary neuron cultures, for example Pax6 and Sox2 as stem cell maintenance factors, and Neurogenin2 as a marker of specified neuronal fate (Figure 7P-R).

While Pax6 levels decreased during WT culture maturation, Pax6 levels remained stable in Nedd8-deficient neurons (Figure 7N). Regarding Ngn2, its levels increased during culture maturation, but not in Nedd8-depleted neurons (Figure 7O). Finally, Sox2 levels remained low in WT conditions, whereas Sox2 levels increased during culture maturation in Nedd8-KO neurons (Figure 7P). This last result contrasts with the RNA-seq result obtained in DIV12 neurons, which indicate similar levels of *Sox2* transcript in both control and Nedd8-depleted neurons (7N), indicating that the Sox2 protein levels are aberrantly regulated in Nedd8-KO neurons. Taken together, the failure to maintain the adequate protein expression levels of key regulators of glutamatergic cell fate in Nedd8-deficient neurons likely explains the aberrant expression of synaptic genes required for the final expression of a mature glutamatergic identity and the aberrant synaptic transmission.

## Discussion

PTMs are essential throughout the lifetime of a nerve cell, from early development and differentiation to synaptic signaling and plasticity. The present study focused on neddylation, an enigmatic, Ubl-based PTM that has recently attracted major attention due to studies linking neddylation to neurite outgrowth, spine growth, synapse density, and synaptic transmission (Vogl et al. 2015; Vogl et al. 2020; Scudder and Patrick 2015; Brockmann et al. 2019; Kang et al. 2021; Song et al. 2023). For optimal stringency in analysing effects of perturbed neddylation, we employed a conditional Nedd8-KO mouse line and studied neuronal development and synaptic transmission in hippocampal autaptic neurons depleted from Nedd8 via infection with CRE-expressing virus, RFP-expressing virus or no viral infection were used as controls. We found that Nedd8-deficient neurons (i) have a moderately reduced dendrite complexity but normal numbers of synapses (Figure 1), (ii) exhibit an increase in synaptic release probability, partly due to altered coupling between SVs in the RRP and VGCCs (Figure 2 and 4), accompanied by a reduced paired-pulse ratio and more profound synaptic depression (Figure 2H-2L) and (iii) show no change in the RRP refiling rate normalized to the RRP_40Hz_ size (Figure 2N) or in VGCC function (Figure 3).

On the other hand, Nedd8 deletion does not cause apparent postsynaptic defects. Some of these results, particularly the observation that postsynaptic features are unaffected by Nedd8-loss, are at odds with previous findings (Vogl et al. 2015; Brockmann et al. 2019; Scudder and Patrick 2015). This is likely due to the different neuron culture models used and the different methodology employed to perturb neddylation. For instance, we used a genetic Nedd8-KO strategy to abolish neddylation while others based their studies on the pharmacological perturbation of neddylation using the Nae1 inhibitor MLN-4924, which has known off-target effects (Mao and Sun 2020; Zhang et al. 2022).

### Control of transmitter release probability by Nedd8, vGluts, and endophilin 1

Our transcriptome analysis revealed that Nedd8-loss leads to a down-regulation of many genes encoding synaptic proteins. A strong decrease in vGlut1 and an increase in vGlut2 expression levels stood out (Figure 5, Table 1). This is consistent with previous studies linking reduced vGlut1 levels to neddylation defects (Vogl et al. 2015; Kang et al. 2021; Song et al. 2023), and resonates well with the correlation of synaptic vGlut1 and vGlut2 levels with respective lower and higher transmitter release probability (Weston et al. 2011; Wojcik et al. 2004). While the total number of synapses was not changed in Nedd8-KO neurons, the 60-70% reduction in vGlut1 levels (Figure 5) was paralleled by a substantial decrease in the number of vGlut1-containing synapses, without changes in the subsynaptic distribution of the remaining vGlut1 (Figure 6). The vGlut1-loss in Nedd8-KO neurons is offset by a 50% increase in vGlut2 levels (Figure 5), which likely compensates for the loss of vGlut1, explaining why mEPSC amplitudes, which are co-determined by SV filling, are not affected by Nedd8-loss (Figure 2).

In view of the fact that high synaptic vGlut2 levels are linked to increased P_vr_ (Wojcik et al. 2004; Weston et al. 2011), we tested the hypothesis that the increased vGlut2-vGlut1 ratio might be at the basis of the increased P_vr_ in Nedd8-KO neurons. This hypothesis was favored because VGCC function was unchanged in Nedd8-KO neurons (Figure 3), and only slight changes in SV-VGCC coupling were observed (Figure 4). However, neither reexpression of vGlut1 nor knock-down of vGlut2 in Nedd8-KO cells reverted the altered P_vr_ (Figure 7). We currently attribute this to the reduced endophilin 1 levels in Nedd8-KO neurons, which are not reverted by vGlut1 re-expression or vGlut2 knock-down (Figure 7), because the effect of vGlut1 expression levels on the P_vr_ is known to be dependent on the presence of endophilin 1 (Weston et al. 2011; Voglmaier et al. 2006). In accord with this notion are the facts that the endocytotic regulator endophilin 1 operates in release site clearance (Haucke, Neher, and Sigrist 2011), and has further been linked to RRP maintenance and the support of sustained exocytosis (Gowrisankaran et al. 2020), which are compromised in Nedd8-KO neurons (Figure 2).

### Profound transcriptome changes upon Nedd8-deletion

To obtain deeper insight into the molecular mechanisms that cause the deficits in Nedd8-KO neurons, we analyzed the neuronal transcriptome upon Nedd8-deletion. This approach was chosen to directly evaluate the impact of Nedd8-loss on the neuronal transcriptional landscape and to infer signaling events that can be linked to the physiological changes we have observed (Roig Adam et al. 2023).

RNA-seq and differential expression analysis revealed an unexpected consequence of Nedd8-loss, characterized by a strong decrease in the expression of genes encoding synaptic proteins, especially when considering the overall upregulation of the rest of the transcriptome. After stringent filtering, 77% of all transcriptomic changes reflected an upregulation of gene expression upon Nedd8-loss (Figure 5A). The subset of affected SynGO genes behaved very differently, with half upregulated and the other half downregulated, indicating that Nedd8-deletion strongly reduces the expression of genes encoding synaptic proteins, while the rest of the transcriptome is mainly upregulated.

Gene ontology and pathway analysis showed that upregulated genes are associated with cancer (Supplementary Figure 2 and 3). The molecular functions, cellular components, and biological processes associated with genes upregulated upon Nedd8-depletion represent a broad range of pathways that are not necessarily interconnected (Supplementary Figure 2). In contrast, genes downregulated upon Nedd8-depletion represented a more restricted functional spectrum, with glutamatergic synapses, glutamatergic signaling, neuronal development, synaptic vesicle cycle, and synaptic plasticity featuring prominently (Supplementary Figure 3). In particular, we found an imbalance in the vGlut1-vGlut2 expression ratio, which reflects the maturation of glutamatergic neurons. As neuronal morphology was only moderately affected while glutamatergic synaptic features showed strong changes upon Nedd8-loss, it appears that blocking neddylation impacts glutamatergic neurons quite specifically rather than neurodevelopment in general. Overall, our data reveal a previously unkown role of neddylation in regulating the transcriptional landscape in neurons, with a strong influence on genes involved in excitatory synaptic signaling.

### Neddylation and neuronal differentiation

Given the substantial transcriptome changes caused by Nedd8-depeletion, the alterations we observed in Nedd8-KO neurons, such as the morphological changes and the increase in transmitter release probability, are likely the final manifestations of a defined but complex remodeling of the transcriptional landscape. This, in turn, is likely linked to the function of cullins, which are the predominant Nedd8 substrates, part of the major class of Cullin-RING E3 ubiquitin ligases, and involved in a broad range of biological processes, including cell growth, development, signal transduction, transcriptional control, and genome integrity (Sarikas, Hartmann, and Pan 2011; Lin and Komives 2024).

As mentioned above, the differentiation of glutamatergic neurons entails a switch from vGlut2 to vGlut1 expression as the neurons mature (Boulland et al. 2004). In view of this, the increased vGlut2-vGlut1 ratio we observed in glutamatergic Nedd8-KO neurons might reflect a delay in the differentiation and maturation of these cells. Indeed, our transcriptome analysis revealed elevated levels of transcription factors found in neural progenitors (Figure 7N, O, Pax6, Sox2), and decreased levels of markers for neural commitment (Figure 7N, O, NeuroD1, NeuroD2, NeuroG2, Trb1) (Hodge, Kahoud, and Hevner 2012; Mihalas and Hevner 2017). Strikingly, Sox2, a key transcriptional regulator involved in stem cell maintenance, is normally degraded during neural progenitor cell differentiation by a pathway that involves Cullin-RING-E3-ligases (Cui et al. 2018). Thus, failure to downregulate Sox2 levels during neural differentiation delays the subsequent increase in expression of transcriptional regulators such as Neurogenin2 and NeuroD1 in Nedd8-KO neurons (Kuwabara et al. 2009), thereby delaying the developmental program of glutamatergic neurons (Figure 7N). In this context, it is important to note that we eliminated Nedd8 expression in P0 mouse neurons cultured for one day. At this stage, neurons have excited the cell cycle and are committed, so that neurogenesis and basic morphogenesis are hardly affected while the expression of the glutamatergic cell identity, which continues postnatally, is.

In general accord with the notion above, neddylation was reported to contribute to the regulation of neurogenesis and cell fate identity. For instance, Nae1 ablation in neuronal progenitor cells leads to severe morphological defects, likely due to impaired neurogenesis and neuronal differentiation (Zhang et al. 2020). Furthermore, Nae1 deletion in Schwann cells causes severe alterations in differentiation pathways, ultimately leading to peripheral neuropathies (Ayuso-Garcia et al. 2024).

### Conclusion

Neddylation is a PTM that affects a plethora of protein targets and cellular processes. The present study shows that neddylation plays a key role in the differentiation and function of glutamatergic neurons. Supporting this notion, our data indicate that Nedd8-ablation in young, postmitotic, glutamatergic neurons causes alterations in the coordinated expression, activation, or homoeostasis of developmental transcription factors that control neuronal differentiation, leading to aberrant transcriptional control and a defects in the development of a mature glutamatergic neuronal phenotype. This becomes manifest in multiple ways, including increased vGlut2 expression level, reduced vGlut1 and endophilin 1 expression levels, reduced dendrite complexity, and an increased probability of neurotransmitter release, partly due to altered coupling between SVs in the RRP and VGCCs.

## Methods

### Materials availability

Material described in this study is available upon request. This includes antibodies, DNA constructs or mouse lines.

### Data and code availability

RNA-seq data are accessible under the GSE269898 and are publicly available as of the date of publication.

Data described in this study, such as Western blot or microscopy data, will be shared upon request by the lead contact. Any additional information required to reanalyze the data reported in this work is available from the lead contact upon request.

This paper does not report original code. Jupiter Notebook for graph generation can be shared upon request.

## Experimental model and details

### Animals

#### Generating *Nedd8* mouse mutant using CRISPR/Cas9 gene editing

The Nedd8cKO knock-out mouse line was generated by site-directed CRISPR-Cas9 mutagenesis. Superovulated FVB/N females were mated with FVB/N males and fertilized eggs collected. In-house prepared CRISPR reagents (hCas9_mRNA, sgRNAs, preformed Cas9_sgRNA RNP complexes, and the dsDNA (HDR fragment) used as a repair template with the required two loxP insertions), were microinjected into the pronucleus and the cytoplasm of zygotes at the pronuclear stage using an Eppendorf Femtojet. Importantly, all nucleotide-based CRISPR-Cas9 reagents (sgRNAs and hCAS9_mRNA) were used as RNA molecules and were not plasmid-encoded, reducing the probability of off-target effects, due to the short live of RNA-based reagents (Doench et al. 2016; Tycko et al. 2019). The sgRNAs targeting the Nedd8 Intron1 (201bp upstream Exon2) and Intron3 (127bp downstream Exon3) were selected using the guide RNA selection tool CRISPOR (Haeussler et al. 2016; Concordet and Haeussler 2018). The correct site-specific insertion of the HDR fragment was confirmed by localization PCR with primers upstream and downstream of the HDR sequence, followed by cloning of the obtained PCR products and sequencing of the obtained clones.

Mice were bred in the animal facility of the Max Planck Institute for Multidisciplinary Sciences (MPI-NAT, City Campus). Nedd8cKOs were classified as unburdened and showed no signs of pathogens, according to standard ethical guidelines.

Mouse maintenance and breeding were performed with permission of the Niedersächsisches Landesamt für Verbraucherschutz und Lebensmittelsicherheit (LAVES). Animals were kept in ventilated cages (TYPE III, 800 cm2) at 21±1°C, 55% relative humidity and under a 12h/12h light dark cycle in specific pathogen-free conditions. Cages were changed once a week. Animals received *ad libitum* access to autoclaved food and water and were provided with bedding and nesting material. Animals’ health was controlled daily by caretakers and a veterinarian. Health monitoring was done quarterly according to FELASA recommendations with NMRI sentinels or animals from the respective colony. The sex of neonatal animals used for experiments was not checked.

#### Nedd8 sgRNAs

Nedd8-protospacer-sgRNA1 sequence (upstream Nedd8_Exon2): 5’-AACCTCAGTCCACAAGCATG-3’(PAM = GGG)

Nedd8-protospacer-sgRNA2 sequence (downstream Exon3) 5’-CTGCGAAAGAGCGGCAAGCA-3’(PAM = AGG)

**Nedd8-Exon2-Exon3-floxed_HDR fragment, 2891bp:**

**CDS Exon2, Exon3 & Stop-Exon4 = BLACK BOLD UPPER CASE**

**loxP1 & loxP2 = BLUE BOLD UPPER CASE**

**STOP Codon TGA = RED BOLD UPPER CASE**

**3’-UTR** = <u>underlined</u>

5’-taagagcacataccgaagccgttgggcacactcctaaaatcacagcagttagagaaagctgagacaggaaaattacaagggtgagttccaggccagtctaggctgcctagtcaaagcaatacaaaagatgatgggtggattagaggttgtgtgaatcacagatttacagtttgtacacaaatatatttggttacctagttttagcctgtggaaatcaagcaatttgagttggagtttaggatttagatagaattctgggacactaggaaagaatataatgttcaactctagcatttcaaaaaaaatctagttggtggtgattacttgtgcccttgaaaactctttgcattatataaatgtaatttgtgccctgggataaagtccttactttaggattgcttaacctagttcagttgtcattgcatctcatccaaagaattcctgatattcttgctgacttggggtgggggtggggaacggcttgtggcattaaggggatcaaatgggttcgcagaatgctgtcttacactgcctgatgacatgggctgtggcacagccttctgattcttcagggtagtattgctgagaatatatacataacaactctcagcagagagaaccccagagaccaagataagatgttacacttagaacatgtcttgccaggcaatggtagtgcatacctttaatcccagcacttggaaggcaaagatcagggccacacaaaataaaccctgtcaccaaaaaaaaaaaattcttctatttttgtgaaaagacactatgataaaagaacgtatttaatttggagcttaccgtttcagaggggaagtccaatactatcatggcagaga ggatggcagcagacaggtaggcatggttctggagcagaagctaagagcttaaacctcagtccaca**ATAACTTCGTAT AATGTATGCTATACGAAGTTAT**agcatgaggcagagaaagagaacatggcttgggcttttgaagcctccaaca agaccacacctctaatacttcccgttccaccaactgtgtgccatgtattcaaacatgagcctattctcattcagtccaccacagaa agcaccttggtatcgggctgtgggacttacgccatctttgcctgttttag**ACGCTGACTGGGAAGGAGATTGAG ATAGACATCGAACCCACAGACAAG**gtgagtcaggtgctcagcgtcctttctctgtatgtgtgcttgcatgtgtgtg cttgcatgtgcttatgaccttggtcctttccttctcaggttgtctgtctatacagcctttgttcttagcattggtttgtttgtttgtttgtttattttagttttccgagacagagtttctctctgtgtacccctggctgtcctggaactcagactggccttgaattcatagagagcagcctgtctct accttccaagtgctgggactaaaggctagtatttgtctaattctaagcagcctctgaccacttgtttctccaaag**GTGGAGC GAATCAAGGAGCGTGTGGAAGAAAAAGAAGGGATTCCCCCCCAGCAGCAGCGGCT CATCTACAGTGGCAAGCAAAT**gtaagcttgggtgggaagtgaggtatagctgccttgtgtgaggatgagtgcag gggaaggtgacgaggctacaggttcatcgggcctgtgctgcttcagtgtgtccctgcttg**ATAACTTCGTATAATGTA TGCTATACGAAGTTAT**ccgctctttcgcagtctgatgaagcatatagaatcttctcagaatataagagaaagcataga atattccagacttagagtaaaatagattggattgtaaactaatattgagctagttaacaaactactaaaacaaatttgtagaatagtaatatatgtgcttttatatgcctttaaattgattatgttcaggtatgtatttataggtatatacgcttgagtacaggtgccctcagatttcagagctgtcagatcctcctggaaccttatttaaatgagattgtgagcctcctgaagtgggtactgggacctgaactcaggtcctctgcaagagaagcaatttgagcttttaacttctgagccatctttccagccctccttattaacatcttattagaaggtctaatgattagtctgataattgcttttttcttgtttttacattttcaagatacttgcaaatagtcataatgtgatatcatctaatttctattgtgtaagctatcatattgttaacactgtgggtctctgtattcaaaattaaatactaattacccttagaggttagtaaaagaatatttttctcatacagtttcatagg cctctcagagatccaccaaactcaggttacaaatgcccattttagatggatttctaatgctttttcttctctgtag**GAATGATGA GAAGACAGCAGCTGATTACAAGATTCTAGGTGGTTCCGTCCTCCACCTGGTGTTGGC TCTTAGAGGAGGAGGTGGTCTTGGGCAGTGA**agaaacttggttccgtttacctccttgccctgccaatcat aatgtggcatcacatatcctctcactctctgggacaccagagccactgccccctctcttggatgcccaatcttgtgtgtctactggtgggagaatgtgaggaccccagggtgcagtgttcctggcccagatggcccctgctggctattgggttttagtttgcagtcatgtgtgcttccctgtcttatggctgtatccttggttatcaataaaatatttcctggccatctggactctttcttcttgacacaaataactgaaatccaagtggctataaccagtagaagtagaacggtggaaggaaaggtgctgctcttaggttagttttgagaaggtgtgggtgggata gg-3’

### Gentotyping Strategie

For genotyping, genomic DNA (gDNA) was isolated from tail biopsies using a genomic DNA isolation kit (Nexttec, #10.924).

For the location PCR, 20 μL reactions were prepared using 1 μL clean gDNA (15-80ng), 4 µL PrimerSet (4 pmol final each), 4 µL 5X Reaction Buffer (Finnzymes #F-524), 0.4 µL PhireHot-Start II Taq DNA Polymerase (Finnzymes #F-122L), 1 µL 10 mM dNTPs (Bioline #DM-515107), 4 µL Hi-Spec Additive (Bioline #HS-014101) and 5.6 µL H2O. Thermocycler parameters: 98°C for 5 min, (98 °C for 45 s, 64 °C for 30 s, 72 °C for 60 s) repeated for 34 cycles, final step at 72 °C for 10 min.

For the diagnostic routine genotyping PCR, 20 μL reactions were prepared using 1 μL clean gDNA (15-80ng), 4 µL PrimerSet (4 pmol final each), 4 µL 5X Reaction Buffer (Biozym #331620XL), 0.2 µL Hot-Start Taq DNA Polymerase (Biozym #331620XL), 1 µL 50 mM MgCl2 (AGCTLab stock) and 9.8 µL H2O. Thermocycler parameters: 96°C for 3 min, (94 °C for 30 s, 62 °C for 60 s, 72 °C for 60 s) repeated for 32 cycles, final step at 72 °C for 7 min.

Primer sequence and fragment size information: Location PCR to confirm the site-specific insertion Primers information:

38438 5’-CTAGGCCAAGTGTGGTGTTTGTGGT-3’

s_Up_HDR1_Nedd8-cKO location PCR

38445 5’-AATCCCAGCAGACAAATCCATGTGA-3’

as_Down_HDR1_Nedd8-cKO location PCR

Fragment sizes:

Nedd8_WT = 2983bp (38438_38445_2983bp)

Nedd8_cKO_floxed = 3051bp (38438_38445_3051bp)

Routine genotyping diagnostic PCR:

Primers information:

38447 5’-CGTATTTAATTTGGAGCTTACCGTT-3’

Sense2_Nedd8_UpExon2

38448 5’-GCAAAGATGGCGTAAGTCCCACA-3’

Asense3_Nedd8_UpExon2

38450 5’-GGCACCTGTACTCAAGCGTA-3’

Asense_Nedd8_DownExon3

Fragment sizes:

Nedd8_UpEx2_WT = 302bp (38447_38448_302bp

Nedd8_cKO_LoxP1 = 336bp (38447_38448_336bp)

Nedd8_KO = 388bp (38447_38450_388bp), after Cre recombination

### Cell lines

HEK293-FT were kept in DMEM medium (Gibco) containing 10% FBS and antibiotics (Gibco, 100 U/ml Penicillin, 100 μg/ml streptomycin, and 500 µg/ml gentamycin) at 37°C with 5% CO_2_.

### Primary hippocampal neuron culture

Continental cultures of primary hippocampal neurons were performed as described (Ripamonti et al. 2020a). Hippocampi were dissected from homozygous Nedd8cKO P0 mice. The tissue was digested in a solution containing 1 mM CaCl_2_, 0.5 mM EDTA, 1.65 mM l-cysteine in DMEM (Gibco) for 1 h at 37°C under agitation (450 rpm). Digestion was stopped via a 15 min incubation at 37°C in DMEM supplemented with 10% heat-inactivated fetal bovine serum (FBS, Gibco), 2.5 mg albumin, and 2.5 mg/ml trypsin inhibitor. The tissue was washed twice with NBA medium containing 500 µL Neurobasal A medium (NBA, Gibco), 1% B27 (Gibco) and 20 µg/ml Penicillin-Streptomycin (Gibco). Careful trituration was performed manually in 100 µL of the same medium. Once the tissue was dissociated, the supernatant was plated onto 60 mm Petri dishes (CytoOne) previously coated with Poly-L-Lysine (Sigma) at an approximat density of 200.000 cells/well (two hippocampi). Medium was replaced the next day. Cultures were kept at 37°C with 5% CO_2_.

### Primary autaptic culture

Primary autaptic neuronal cultures and astrocyte micro-island cultures were performed as previously described (Burgalossi et al. 2012). Briefly, hippocampi were dissected from homozygous Nedd8cKO P0 animals. Tissue digestion, washing and trituration steps were performed as described above for primary continental cultures. Neurons were plated on 6-well astrocyte micro-island plates at a density of 4.000 cells/well. Plates were kept in NBA media containing Neurobasal A medium (Gibco), 1% B27 (Gibco) and 20 µg/ml Penicillin-Streptomycin (Gibco) at 37°C and 5% CO_2_.

### Animals Lentivirus and AAV production

Lentivirus production was performed as previously described (Follenzi and Naldini 2002b, 2002a; Lois et al. 2002; Salmon and Trono 2006). In brief, HEK293-FT (Thermo) cells grown in DMEM medium containing 10% FBS and antibiotics (100 U/ml Penicillin, 100 μg/ml streptomycin, and 500 µg/ml gentamycin) with an 80-90% confluency were transfected with 20 µg of pVSVG packaging and pCMV delta R8.9 envelope constructs, and 40 µg of lentivirus construct expressing under the Synapsin1 promoter under the Synapsin1 promoter either red fluorescent protein (RFP) or CRE-recombinase tagged with an RFP protein (CRE). Transfection was performed using Lipofectamine 2000 (Thermo Scientific). After 6h, medium was changed to Optimem (Gibco) supplemented with 2% FBS, penicillin 100 U/ml, Streptomycin 100 U/ml, and 10 mM sodium butyrate. After 46-48h, the cell supernatant was centrifuged at 2.000 rpm for 5 min at 4°C and then filtered using a 0.45 µm filter (Merck). Viral particles were filtered and washed with Optimem once and twice with 1x tris-buffered saline (TBS) using 100 kDa Amicon centrifugal filter (Millipore) via series of centrifugation at 3.500 rpm for 10 min at 4°C. Viral particles were concentrated to a final volume of 500 µL and aliquots of 50 µL were snap frozen in liquid nitrogen. vGlut1 and vGlut2 rescue viral constructs, as well as empty control constructs, were provided by the Viral Core Facility of Charité Universitätsmedizin Berlin.

### Electrophysiology

Whole-cell patch-clamp recordings were performed in autaptic excitatory hippocampal neurons at DIV10-12. All recordings were done using a HEKA EPC-9USB amplifier controlled by PatchMaster software (HEKA electronics), the recording rate was 10 kHz, and R_series_ up to 10 MΩ with 50% compensation. The membrane potential of excitatory neurons was set to -70 mV in voltage-clamp configuration and depolarized to 0 mV to induce action potentials. For synaptic properties, the extracellular solution contained (mM); 140 NaCl, 2.4 KCl, 10 HEPES, 10 Glucose, 4 CaCl_2_, 4 MgCl_2_, pH 7.6. Microelectrodes were fabricated from borosilicate glass pipettes using a micropipette puller (P-27, Sutter Instrument) and filled with intracellular solution containing (mM) 136 KCl, 17.8 HEPES, 1 EGTA, 4.6 MgCl_2_, 4 NaATP, 0.3 Na_2_GTP, 15 creatine phosphate, and 5 U/ml phosphocreatine kinase osmolarity 315-320 mOsmol/L, pH 7.4. Pipette resistances were around ∼ 2-4 MΩ. Excitatory post-synaptic currents (EPSC) were recorded using frequencies of 0.2 Hz, 10 Hz, and 40 Hz. The release of vesicles from the readily releasable pool (RRP) was triggered by application of a hypertonic 0.5 M sucrose solution. Estimation of the size of the RRP was obtained from the hypertonic sucrose treatment and back extrapolation of the cumulative EPSC from 40 Hz trains of action potential (Neher 2015). Miniature EPSC (mEPSC) were recorded under 300 mM tetrodotoxin (TTX, Tocris) treatment, and for post-synaptic responses, 100 µM Glutamate (Sigma) or 3 µM γ-aminobutyric acid (GABA) (Sigma) solutions were used. The paired-pulse ratio was calculated from the 40 Hz train stimulus and is the ratio of the peak amplitudes of the second and first responses. For calcium channel sensitivity studies, the extracellular solution contained (mM) 140 TEA-Cl, 2.5 CsCl, 10 CaCl_2_, 1 MgCl_2_, 10 HEPES, 10 glucose, Osmolarity 310 mOsm, pH 7.3; and the intracellular solution contained 130 CsCl, 10 HEPES, 2 CaCl_2_, 10 EGTA, 5 MgATP, Ph 7.3. 10 µM Nimodipine (Alomone Labs) or 0.2 µM ω-Agatoxin (Alomone Labs) (L-type and PQ-type calcium channel blocker, respectively) were used to block specific calcium channels. For synaptic vesicle coupling studies, 100 µM EGTA-AM (Invitrogen) was used. Infected neurons were distinguished by nuclear RFP signal that was visualized using an inverted microscope (Zeiss). Data was analyzed using AxoGraph software (Axograph Scientific).

### Western blot

Neuronal cultured were lysed in lysis buffer containing 20 mM Tris Base, 150 mM NaCl, 1% Triton X-100 (Roche), and protease inhibitors (0.5 µg/ml aprotinine, 0.2 mM phenylmethylsulfonyl fluoride, 1 µg/ml leu-peptine, 20 mM N-ethylmaleimide) pH 7.4-7.6. For storage, samples were snap-frozen in liquid nitrogen and kept at -80°C. SDS-PAGE was performed using self-made 10% acrylamide gels, or commercial, pre-casted 4-12% Bis-Tris gels (Thermo Scientific). Samples were sonicated and protein concentration was determined using Bradford assay (Biorad). Samples were heated for 5 min at 95°C, or for 20 min 65°C, shortly before loading. Gel migration was performed in a buffer containing 5 mM MOPS, 5 mM Tris Base, 0.01% SDS, 0.1 mM EDTA, pH 7.7. Proteins were transferred to nitrocellulose membranes (Cytiva) using a buffer containing 192 mM Glycine, 25 mM Tris Base, and 20% Methanol. Total protein content was visualized using the Memcode reversible protein staining kit (Thermo Scientific). Primary and secondary antibody incubation of the membrane were performed in 1x PBS supplemented with 5% milk (Frema) and 0.1% Tween 20 (PBS-T) following standard procedure. Signal was revealed using enhanced chemiluminescent (GE Healthcare) and signals were detected with an ECL Chemostar Imager (Intas Science Imaging) and analyzed with Fiji (NIH). Quantification was performed by normalizing the signal in each lane with its respective Memcode signal.

### RNA isolation, RNA-sequencing, and analysis

Continental culture of primary hippocampal neurons aged for 13 days were subjected to TRIzol-RNA extraction following manufacture protocol (Zymo). RNA quality was assessed by measuring the RNA integrity number (RIN) using a Fragment Analyzer HS Total RNA Kit (Advanced Analytical Technologies, Inc.). Library preparation for RNA-Seq was performed in the STAR Hamilton NGS automation using the Illumina Stranded mRNA Prep, Ligation (Cat. N°20040534 and the Illumina RNA UD Indexes Set A, Ligation, 96 Indexes, 96 Samples Cat. N°20091655) starting from 300 ng of total RNA. The size range of the final cDNA libraries was determined by applying the SS NGS Fragment 1-to 6000-bp Kit on the Fragment Analyzer (average 340 bp). Accurate quantification of cDNA libraries was performed by using the QuantiFluor™ dsDNA System (Promega). cDNA libraries were amplified and sequenced by using an S4 flow cell NovaSeq6000; 300 cycles, 25 Mio reads/sample from Illumina.

#### Raw read & Quality check

Sequence images were transformed with BaseCaller Illumine Software to BCL files and demultiplexed to fast files with bcl2fastq v2.20.0.422. The sequencing quality was asserted using FastQC v.0.11.5 software (http://www.bioinformatics.babraham.ac.uk/projects/fastqc/) (Wingett and Andrews 2018).

#### Mapping and Normalization

Sequences were aligned to the reference genome Mus musculus (GRCm39 version 110, https://www.ensembl.org/Mus_musculus/Info/Index) using the RNA-Seq alignment tool (Dobin et al. 2013)(version 2.7.8), allowing for 2 mismatches within 50 bases. Subsequently, read counting was performed using featureCounts (Liao, Smyth, and Shi 2014). Read counts were normalized in the R/Bioconductor environment (version 4.3.1) using the DESeq2 (Love, Huber, and Anders 2014) package version 1.40.2.

#### PCA and other quality checks

Principal component analysis (PCA) was performed using sklearn Python library (version 1.3.1). [Pedregosa F, Varoquaux G, Gramfort A, Michel V, Thirion B, Grisel O, Blondel M, Müller A, Nothman J, Louppe G et al. Scikit-learn: machine learning in python. ArXiv 2012, abs/1201.0490.] (Supplementary Figure 1).

The total gene count box plot displays gene expression levels in each of the samples analyzed (Supplementary Figure 1B). The horizontal line inside each box represents the median gene expression. The box itself encompasses the interquartile range, covering from the 25th to the 75th percentile. The whiskers extend to the 10th percentile on the lower end and the 90th percentile on the higher end. Any points outside the whiskers, particularly those above the 90th percentile, are shown as individual circles. The plot uses color to differentiate between conditions, with red indicating the ‘CRE’ condition and blue representing the ‘RFP’ condition. After PCA and gene count analysis, two samples were removed from the differential analysis: Nedd8Cre1-3 and Nedd8Cre3-3.

#### Differential expression and pathway enrichment analysis

Differential expression analysis was performed using PyDESeq2, a python implementation of DESeq2 method (Love, Huber, and Anders 2014; Muzellec et al. 2023) (version 0.4.4). Pathway enrichment was done with GSEA method (Subramanian et al. 2005) using blitzgsea library (version 1.3.42) (Lachmann, Xie, and Ma’ayan 2022). The calculation of the score in pathway analysis was done as previously described (Hendriks et al. 2018).

### Immunostaining and confocal microscopy

DIV12-13 autaptic hippocampal neurons were fixed with 4% para-formaldehyde (PFA) prepared in 1x phosphate-buffered saline (PBS) for 10 min. All steps were done at room temperature. Neurons were washed three times with 1x PBS and quenched for 10 min with 50 mM Glycine in 1x PBS. Neurons were permeabilized in 1 x PBS containing 0.1% Triton X-100 (Roche), 2.5% goat serum (GS) (Gibco) for 30 min. Cells were washed in 1 x PBS supplemented with 2.5% GS and incubated with primary antibodies diluted in the same solution for 1 h. Neurons were washed and incubated with secondary antibodies for 45 min in darkness. Coverslips were washed in 1x PBS and mounted on slides using Aqua-Poly Mount (Polysciences) medium and stored at 4°C until image acquisition. Images were obtained with a SP8 confocal Leica microscope (Leica Microsystems), 63x immersion objective (NA 1.4), 12-bit image depth, step size of 0.3 µm, and resolution of 1232×1232 corresponding to a pixel size of 149.9 nm. Synapse counting, Sholl analysis and total dendrite lenght were calculated using Imaris software (Version 9.9.1, Oxford Instruments). Representative images were processed using Fiji (NIH).

### STED microscopy and analysis

DIV12 autaptic hippocampal neurons were fixed with 4% PFA for 45 min and then quenched using 100 mM NH4Cl in 1x PBS for 20 min. All steps were done at room temperature. Cells were washed twice in 1 x PBS and incubated in a blocking solution containing 2.5% bovine serum albumin (BSA, Biomol), and 0.1% Triton X-100 (Roche) in 1x PBS. Neurons were incubated with nanobodies against vGlut1-AZD568 (Nanotag), PSD95-STAR635 (Nanotag), and Synaptotagmin1-ATTO488 (Nanotag) in a 1:100 dilution in blocking solution. Coverslips were washed twice in 1x PBS and mounted on slides using Mowiol (Carl Roth). Image acquisition was done using an Abberior Expert line setup (Abberior Instruments) with an IX83 microscope (Olympus), with an 100x immersion objective (1.4 NA). Excitation of STAR635 was performed with a 640 nm laser (5% of max. power) and detected with an avalanche photodiode (APD) laser (range 650-720 nm). For AZD568, excitation was done using a 580 nm laser (30% of max. power) and detected with an APD laser (range 605-625 nm). For both STAR635 and AZD568, depletion was achieved with a 775 nm depletion laser (20% and 60%, respectively). Excitation of ATTO488 was achieved with a 488 nm laser (4% of max. power) and detected with an APD laser (range 525-575 nm). For ATTO488, a solid-state 595 nm depletion laser (10% of max. power) was used. For each neuron, 2-5 regions of interest were selected and imaged using confocal and STED microscopy. Pixel sizes were 200 and 20 nm for confocal and STED, respectively, dwell times 8-10 µs, and lines accumulation of 3-5 for STED. The analysis was performed using Mathlab (The Mathworks, Inc.). Synapses were identified automatically, using band-a-pass filter procedure, which revealed the synapse-sized spots in the different channels. Spots of less than 9 pixels were removed, being considered to be noise events. All synapse images from one experiment were then automatically rotated and oriented, to display the postsynaptic density in the center, and the pre-synapse to the left. Line scans were generated across the synapses, and the intensities in the different channels were measured and displayed. Representative images were processed using Fiji (NIH).

### Statistical analysis

All statistical analyses were performed using GraphPad Prism software (GraphPad). Normality distribution was determined using D’Agostino-Pearson or Shapiro-Wilk normality tests. Statistical analysis was performed using a Kruskal-Wallis and Dunńs multiple comparison test, unpaired t-test, or Mann-Whitney tests. For graphs, data is shown as mean ±SEM. For all figures, statistically significant differences are denoted in graphs with * standing for p-value, where * refers to p<0.05, ** to p<0.01, *** to p<0.001, and **** to p<0.0001. N refers to the number of animals or biological replicate, and n corresponds to the number of cells.

## Acknowledgments

The authors have no conflicting financial interests.

We thank D. Warnecke, and C. Harenberg for their expert technical assistance in DNA synthesis, DNA sequencing, and mouse genotyping, the staff of the Max Planck Institute for Multidisciplinary Sciences Transgenic Animal Facility for the generation and maintenance of mouse colonies, Tanja Nilsson and Sally Wenger for their technical support. RNA-sequencing was performed at the NGS Integrative Genomics (NIG) Core unit of the Institute of Pathology at rthe University Medical Center Goettingen. We thank Thorsten Trimbuch and Bettina Brokowski from the viral core facility of the Charité – Universitätsmedizin Berlin for providing vGlut1 and vGlut2 rescue constructs.

This work was supported by the German Research Foundation (SFB1286/A9, MT, NB).

## Author contribution

MT designed the projects, performed experiments and wrote the manuscript with the help of all other co-authors. JT performed experiments. NB, JSR, CR, FB, SR provided material, conceptual feedback and edited the manuscript. MF and SB analyzed the RNA-seq dataset.

## Supplemental Information

**Supplementary Figure 1:**
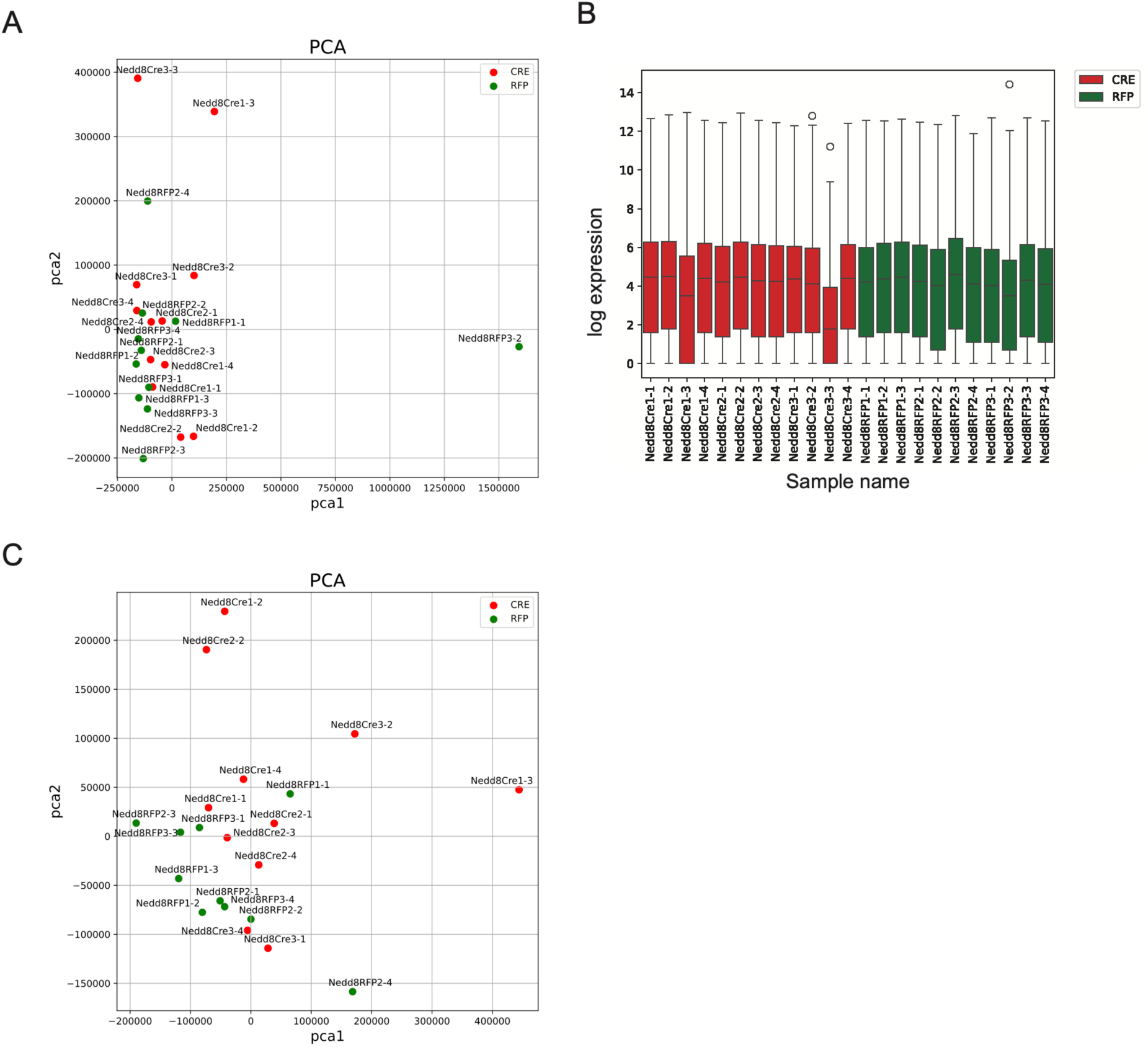
Quality check for the RNA-seq: PCA and total gene counts.

**Supplementary Figure 2:**
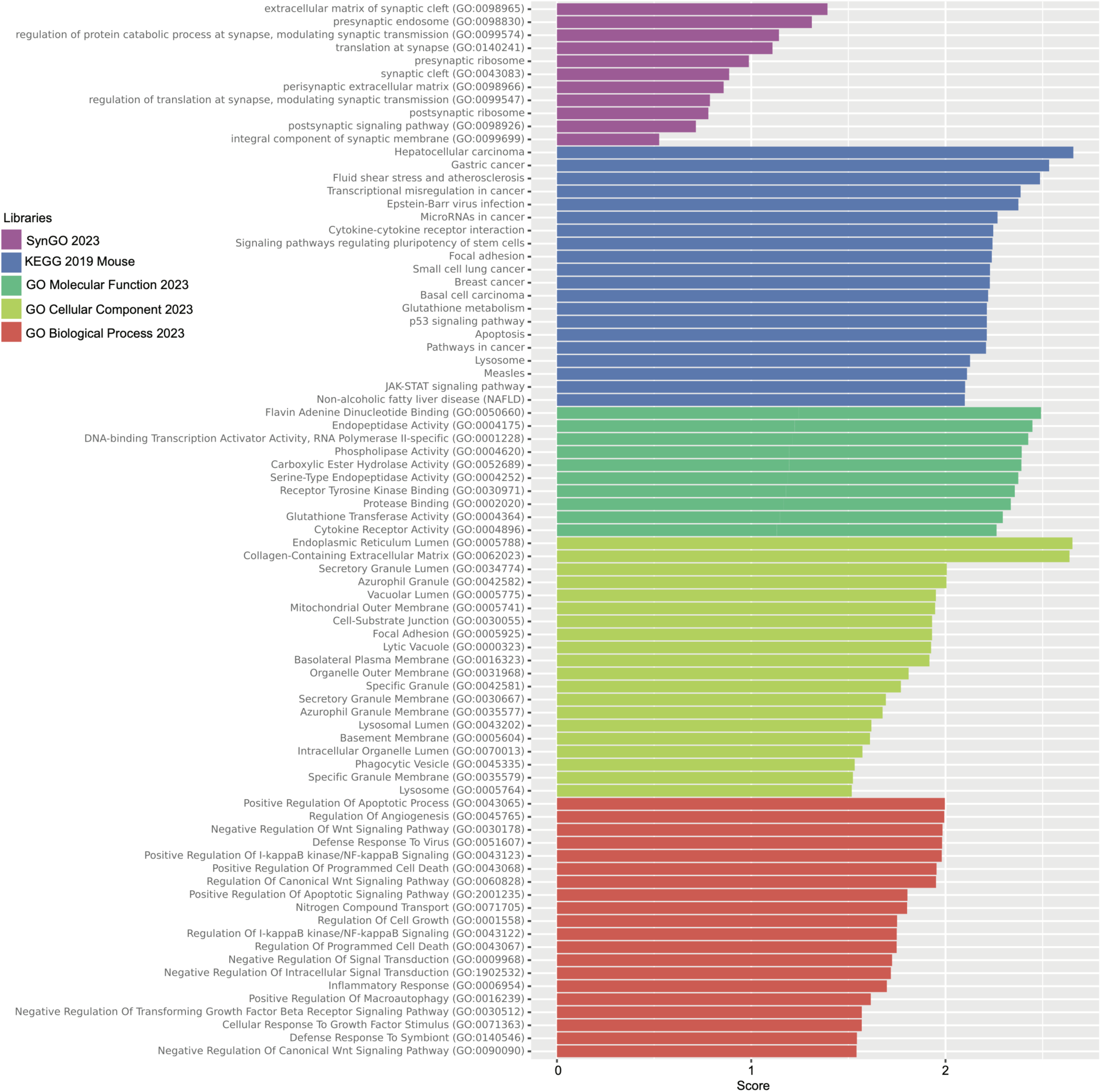
Gene ontology of up-regulated genes upon Nedd8 depletion.

**Supplementary Figure 3:**
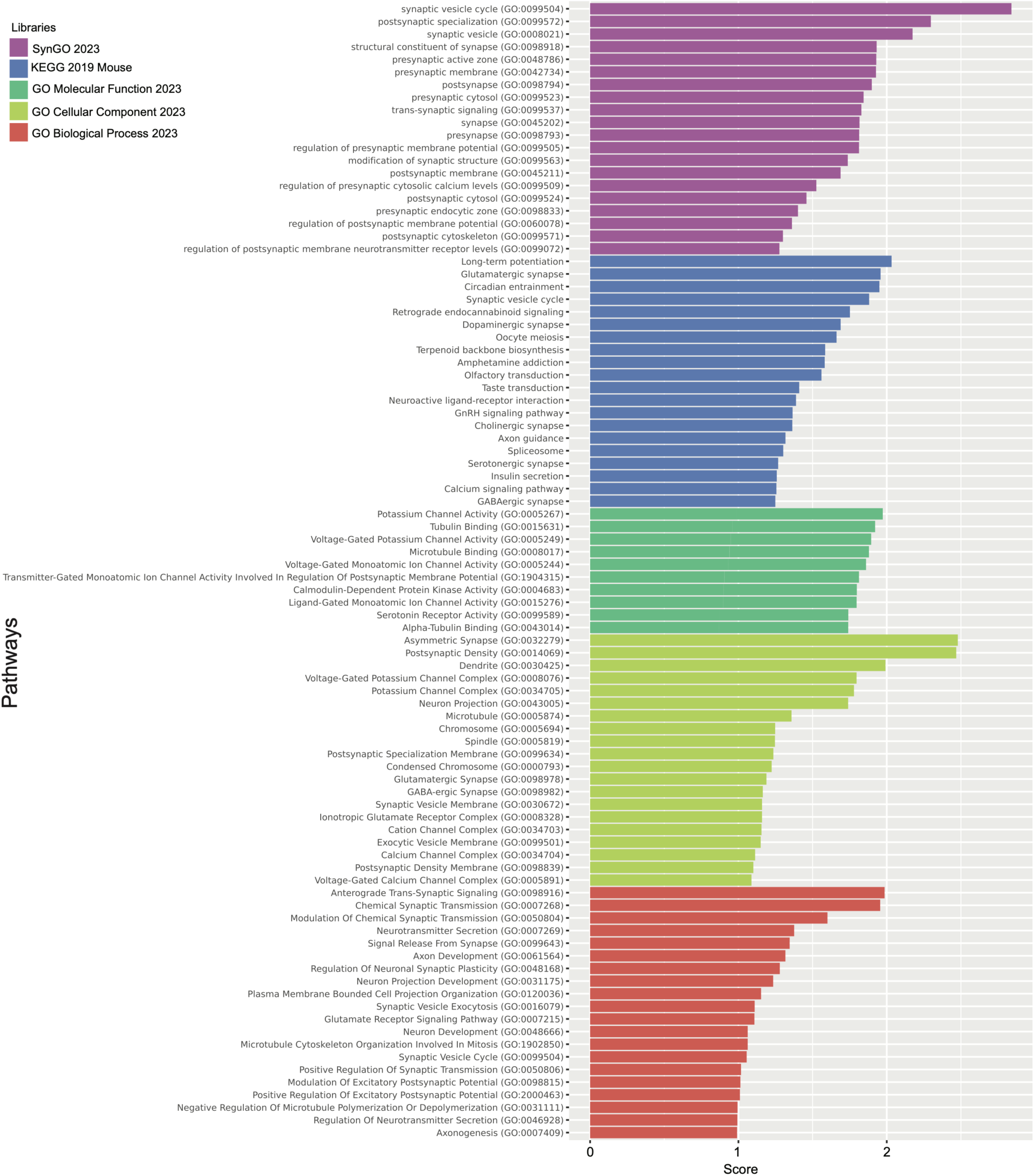
Gene ontology for down-regulated genes upon Nedd8 depletion.

